# Potent SARS-CoV-2 Neutralizing Antibodies Directed Against Spike N-Terminal Domain Target a Single Supersite

**DOI:** 10.1101/2021.01.10.426120

**Authors:** Gabriele Cerutti, Yicheng Guo, Tongqing Zhou, Jason Gorman, Myungjin Lee, Micah Rapp, Eswar R. Reddem, Jian Yu, Fabiana Bahna, Jude Bimela, Yaoxing Huang, Phinikoula S. Katsamba, Lihong Liu, Manoj S. Nair, Reda Rawi, Adam S. Olia, Pengfei Wang, Gwo-Yu Chuang, David D. Ho, Zizhang Sheng, Peter D. Kwong, Lawrence Shapiro

## Abstract

Numerous antibodies that neutralize SARS-CoV-2 have been identified, and these generally target either the receptor-binding domain (RBD) or the N-terminal domain (NTD) of the viral spike. While RBD-directed antibodies have been extensively studied, far less is known about NTD-directed antibodies. Here we report cryo-EM and crystal structures for seven potent NTD-directed neutralizing antibodies in complex with spike or isolated NTD. These structures defined several antibody classes, with at least one observed in multiple convalescent donors. The structures revealed all seven antibodies to target a common surface, bordered by glycans *N*17, *N*74, *N*122, and *N*149. This site – formed primarily by a mobile β-hairpin and several flexible loops – was highly electropositive, located at the periphery of the spike, and the largest glycan-free surface of NTD facing away from the viral membrane. Thus, in contrast to neutralizing RBD-directed antibodies that recognize multiple non-overlapping epitopes, potent NTD-directed neutralizing antibodies target a single supersite.

## Introduction

Severe acute respiratory syndrome coronavirus 2 (SARS-CoV-2), the causative agent for Coronavirus Disease 2019 (COVID-19), emerged in 2019, rapidly establishing an ongoing worldwide pandemic with tens of millions infected and over one million dead (Callaway et al., 2020; Cucinotta and Vanelli, 2020; Dong et al., 2020). In response, an unprecedented global effort to develop vaccines and therapeutics is well underway. One promising approach is the identification of SARS-CoV-2-neutralizing antibodies, which could be used as therapeutic or prophylactic agents. Analysis of such antibodies can reveal viral sites of vulnerability to antibody neutralization, which can help guide the development of vaccines or therapeutics (Burton and Walker, 2020). The primary target for neutralizing antibodies is the viral spike protein, a trimeric type I viral fusion machine (Walls et al., 2020; Wrapp et al., 2020b) that binds virus to the ACE2 receptor on host cells (Benton et al., 2020; Yan et al., 2020; Zhou et al., 2020) and mediates fusion between the viral and cell membranes. The spike protein is comprised of two subunits: the S1 subunit comprising the N-terminal domain (NTD), the receptor-binding domain (RBD) and several other subdomains, and the S2 subunit that mediates virus–cell membrane fusion (Walls et al., 2020; Wrapp et al., 2020b).

The majority of SARS-CoV-2 neutralizing antibodies so far identified target RBD(Brouwer et al., 2020; Cao et al., 2020; Chen et al., 2020; Chi et al., 2020; Ju et al., 2020; Liu et al., 2020b; Pinto et al., 2020a; Robbiani et al., 2020; Rogers et al., 2020; Seydoux et al., 2020; Wang et al., 2020a; Wrapp et al., 2020a; Wu et al., 2020; Zeng et al., 2020; Zost et al., 2020). Structural studies (Barnes et al., 2020a; Barnes et al., 2020b; Liu et al., 2020a; Wang et al., 2020b; Yuan et al., 2020b) and binding competition experiments (Liu et al., 2020a), have revealed neutralizing antibodies to recognize RBD at multiple distinct sites, and further revealed multi-donor RBD-directed antibody classes that appear to be elicited with high frequency in the human population (Barnes et al., 2020b; Robbiani et al., 2020; Wu et al., 2020; Yuan et al., 2020b) as well as in mice with a humanized immune system (Hansen et al., 2020). Neutralization for many RBD-directed antibodies can be explained by interference with RBD-ACE2 interaction, and/or impeding the ability of RBD to adopt the “up” conformation (Barnes et al., 2020b; Liu et al., 2020a; Yuan et al., 2020b) required for ACE2 binding (Benton et al., 2020).

NTD-directed neutralizing antibodies targeting the MERS betacoronavirus have been extensively characterized (Chen et al., 2017; Pallesen et al., 2017; Wang et al., 2018; Zhou et al., 2019). For SARS-CoV-2, a single cryo-EM structure has been reported for the NTD-directed neutralizing antibody 4A8 in complex with SARS-CoV-2 spike (Chi et al., 2020). NTD-directed antibodies have also been observed in electron microscopy (EM) analyses of antibodies from the sera of convalescent donors (Barnes et al., 2020b; Brouwer et al., 2020), and a low-resolution structure of a very potent antibody 4-8 has been reported (Liu et al., 2020a). This report was also notable for the identification of multiple NTD-neutralizing antibodies with potencies rivaling those of the best RBD-directed neutralizing antibodies.

Here we describe cryo-EM and crystal structures for seven potently neutralizing antibodies in complex with either SARS-CoV-2 spike or NTD. We analyzed the genetic basis of recognition for each of the seven antibodies, and further clustered them into antibody classes with similar genetics and modes of recognition. We also analyzed the antibody angles of approach and their recognized epitope. Remarkably, all seven antibodies targeted a single glycan-free surface of NTD, defining an NTD-antigenic supersite. We propose that all potently neutralizing NTD-directed SARS-CoV-2 neutralizing antibodies might target this site.

## Results

### NTD-directed SARS-CoV-2 neutralizing antibodies

Prior studies have identified SARS-CoV-2-neutralizing antibodies that are S1-directed, but do not recognize RBD (Brouwer et al., 2020; Kreer et al., 2020; Rogers et al., 2020; Seydoux et al., 2020; Zost et al., 2020). Other studies have further delineated recognition and shown such antibodies to recognize NTD (Chi et al., 2020; Liu et al., 2020a; Zost et al., 2020). We identified a total of 17 published antibodies, and found that they derived from only 9 VH genes, with antibodies originating from five genes (VH1-24, VH1-69, VH3-30, VH1-8, and VH4-39) evident in multiple donors (**Figure S1**). While these observations were sparse, they raised the possibility that NTD-directed neutralizing responses in different individuals could involve the convergent development of similar antibodies.

Structures for seven such antibodies in complex with SARS-CoV-2 spike or isolated NTD are presented below, grouped by VH gene.

### NTD-directed neutralizing antibodies derived from VH1-24 represent a multi-donor class

Four NTD-directed neutralizing antibodies identified from convalescent donors – antibodies 1-68 and 1-87 derived from donor ‘1’, and 2-51 derived from donor ‘2’ (Liu et al., 2020a) and antibody 4A8 from a third donor (Chi et al., 2020) derive from the VH1-24 gene (**Figure 1A**). In addition to utilizing the same VH gene, three of these antibodies, 1-68, 1-87, and 4A8 utilized an identical set of heavy chain-antibody genes – VH1-24, D6-19, and JH6 – and showed significant similarity in their heavy chain third-complementarity-determining regions (CDR H3s), each of which was 21 amino acids in length (**Figure S2**). Antibody 2-51 also utilized VH1-24, but utilized different D and J genes, D6-13 and JH4, encoding a shorter CDR H3 region of only 14 amino acids. These VH1-24-derived antibodies utilized four different VL-genes, 1-87, 2-51, and 1-69 utilized lambda light chains VL2-14, VL2-8, and VL2-18, respectively, while 4A8 utilized kappa light chain VK2-24.

**Figure 1.**
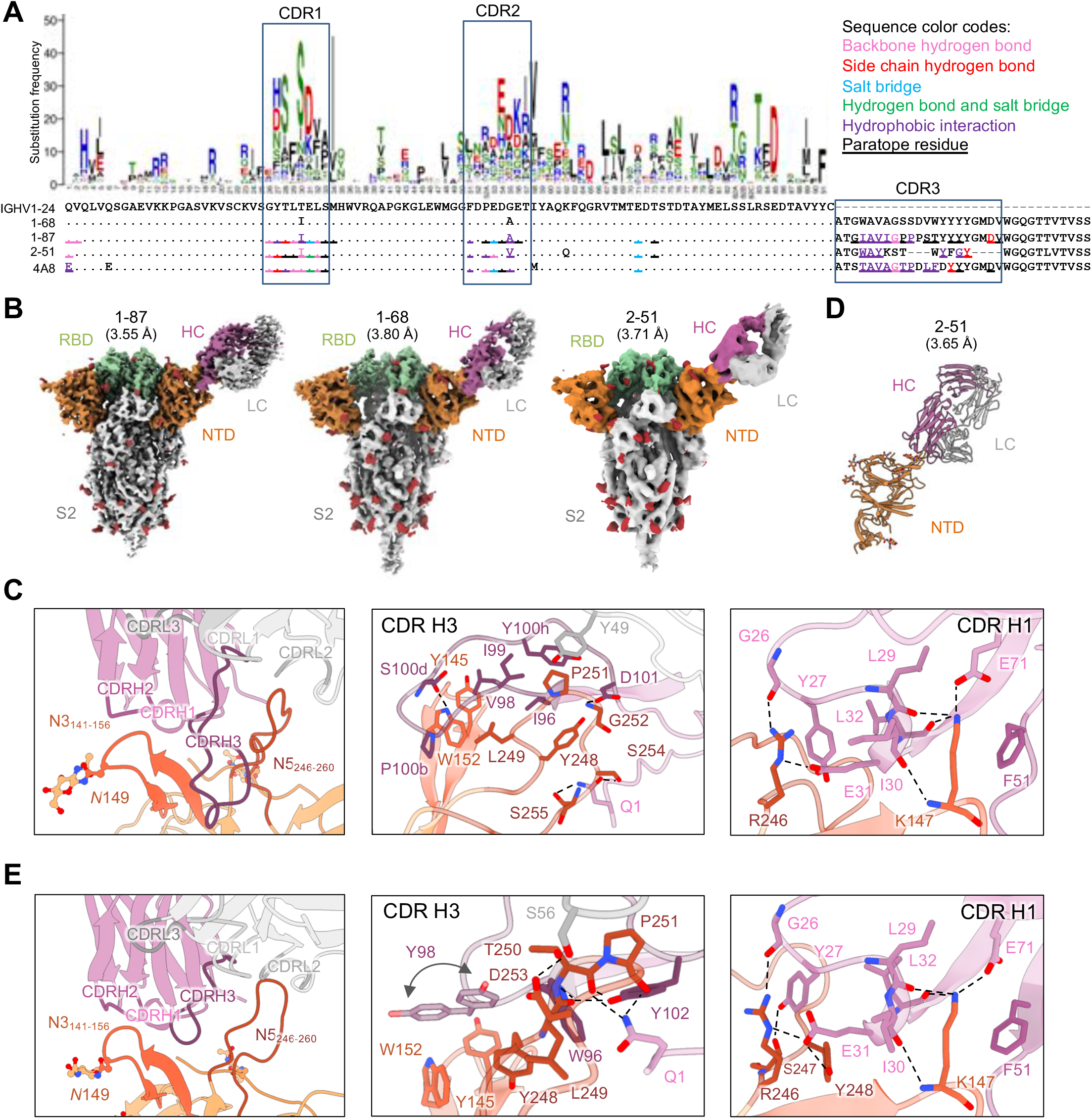
NTD-directed neutralizing antibodies derived from the VH1-24 gene define a multi-donor antibody class. (A) Sequence alignment of VH1-24-derived NTD-directed antibodies showing paratope residues, somatic hypermutations, and gene-specific substitution profile (GSSP) showing somatic hypermutation probabilities for VH1-24 gene. Antibody positions are assigned using the Kabat scheme, the CDRs are assigned by IMGT scheme. Paratope residues are highlighted by underscoring and colored by interaction types. Amino acids in GSSP are colored by chemical property. (B) Cryo-EM reconstructions for spike complexes with antibodies 1-87, 1-68, and 2-51. NTD is shown in orange, RBD in green, glycans in red, with antibody heavy chains in magenta and light chains in gray. (C) Expanded view of 1-87 interactions with NTD showing overall interface (left panel), recognition by CDR H3 (middle panel), and recognition by CDR H1 (right panel). NTD regions N3 (residues 141-156) and N5 (residues 246-260) are colored in shades of orange; CDR H1, H2, H3 are colored in shades of magenta; CDR L1, L2, and L3 are colored in shades of gray. Nitrogen atoms are colored in blue, oxygen atoms in red; hydrogen bonds (distance <3.2 Å) are represented as dashed lines. (D) Crystal structure of antibody 2-51 complexed with NTD, colored as in (B). (E) Expanded view of 2-51 interactions with NTD showing overall interface (left panel), recognition by CDR H3 (middle panel), and recognition by CDR H1 (right panel), colored as in (C). See also Figures S1-S7, Table S1 and Table S2.

We determined cryo-EM structures for the spike complexes with antibodies 1-68, 1-87, and 2-51 at overall resolutions of 3.8 Å, 3.55 Å, and 3.71 Å, respectively (**Figure 1B, Figure S3A and Table S1**). We also produced a locally refined cryo-EM map around the antibody:spike interface for 1-87 at 3.81 Å resolution, which allowed construction and refinement of an atomic model (**Figure 1C**). However, resolution in the antibody:spike interface region was blurred by domain motions for antibodies 2-51 and 1-68. We therefore produced crystals for 2-51 in complex with NTD, which provided an x-ray structure at 3.65 Å resolution (**Figure 1D and Table S2**).

Cryo-EM reconstructions of the VH1-24-derived 1-68, 1-87, and 2-51 antibodies each show a single Fab bound to the NTD of one subunit of the trimeric spike (**Figure 1B**). All target, with similar angle of approach, a single region on NTD – the loop region furthest from the spike-trimer axis. Moreover, the epitope and angle of approach for antibodies 1-68, 1-87 and 2-51 appear similar to those of antibody 4A8 (Chi et al., 2020), also derived from the VH1-24 gene.

Chi et al. (2020) defined the NTD loops in the 4A8-binding region as N1-N5 (corresponding to residue stretches 14 to 26, 67 to 79, 141 to 156, 177 to 186, and 246 to 260, respectively), and we adopt this nomenclature here. We note that the region defined as the N3 loop corresponds to a β-hairpin that includes both β-strands that form a stem region and a short loop that connects them. The N1-N5 loops are disordered in most structures of spike, but some of these loops become ordered in antibody complexes. The structure of antibody 1-87 in complex with spike reveals almost all interactions to be mediated through heavy chain. The 19-residue CDR H3 loop provides the predominant interaction (**Figure 1C, middle panel**), with additional contributions mainly from CDR H1 (**Figure 1C, right panel**). CDR H3, which inserts between the N3 and N5 loops of NTD (**Figure 1C, left panel**) includes several hydrophobic residues (Ile96_HC_, Val98_HC_, Ile99_HC_, Pro100b_HC_, and Tyr100h_HC_) which interact with aromatic residues in N3 including Tyr145_NTD_ and Trp152_NTD_, and with hydrophobic residues of N5. Residues Ser100d_HC_ and Asp101_HC_ in CDR H3 also form hydrogen bonds with Trp152_NTD_ and Gly252_NTD_ respectively; the N-terminal glutamine residue of the heavy chain is also involved in hydrogen bonds with Ser254_NTD_ and Ser255_NTD_ in N5. Residues in CDR H1 form a network of hydrogen bonds involving positively charged residues from N3, Lys147_NTD_, and N5, Arg246_NTD_, with interactions by the side chains CDR H1 residues Tyr27_HC_ and Glu31_HC_. Two additional VH1-24-gene specific glutamic acid residues – Glu53_HC_ in CDR H2 (**Figure S4A**), and framework residue Glu71_HC_ each participate in salt bridges with NTD. The only interaction mediated by the light chain is a hydrophobic interaction between Tyr49_LC_ in CDR L2 and Pro251_NTD_ in N5.

The crystal structure of antibody 2-51 in complex with NTD reveals recognition remarkably similar to that of 1-87. The 14-residue CDR H3 loop of 2-51 inserts between the N3 and N5 loops of NTD (**Figure 1E, left panel**) with additional interactions from CDR H1. CDR H3, includes three aromatic residues (Trp96_HC_, Tyr98_HC_, and Tyr102_HC_), which interact with aromatic residues in N3 including Tyr145_NTD_ and Trp152_NTD_, and with hydrophobic residues of N5 including Tyr248_NTD_, Leu249_NTD_, and Pro251_NTD_ (**Figure 1E, middle panel**). The heavy chain Gln1_HC_ residue is involved in hydrogen bonds with Thr250_NTD_ and Pro251_NTD_ in N5, similar to the hydrogen-bonding pattern observed for 1-87. The only interaction mediated by the light chain is a hydrogen bond between Ser56_LC_ in CDR L2 and Asp253_NTD_ in N5. Residues of CDR H1 form a network of hydrogen bonds nearly identical to the network formed in the 1-87 crystal structure (**Figure 1E, right panel**). Comparison of the 1-87 and 2-51 structures reveals a comparable level of similarity with VH1-24-derived antibody 4A8.

Overall, their common derivation from a common gene and highly similar recognition show the VH1-24 antibodies to define a multi-donor antibody class. Overall, the interaction is dominated by conserved contacts in CDR H1, along with hydrophobic interactions mediated by CDR H3. The VH1-24 gene restriction is explained partly by the conserved interactions of CDR H1 including those of the VH1-24-specific Glu31, and VH1-24-specific residues Glu53 and Glu71 (**Figure S5A**).

### NTD-directed neutralizing antibodies derived from VH3-30 and VH3-33 genes show distinct recognition

Three neutralizing antibodies directed against NTD have been reported that derived from the highly similar VH genes VH3-30 and VH30-33. The high similarity of these genes, which encode only two amino acid differences between them (**Figure 2A**), raised the possibility that these antibodies might represent a multi-donor class despite their derivation from two distinct genes. We therefore determined cryo-EM structures for spike complexes with antibodies derived from each gene: antibody 4-18 from VH3-30 and antibody 5-24 from VH3-33 at 2.97 and 3.9 Å resolutions, respectively **(Figure 2B-C, Figure S3B-C, and Table S1)**.

**Figure 2.**
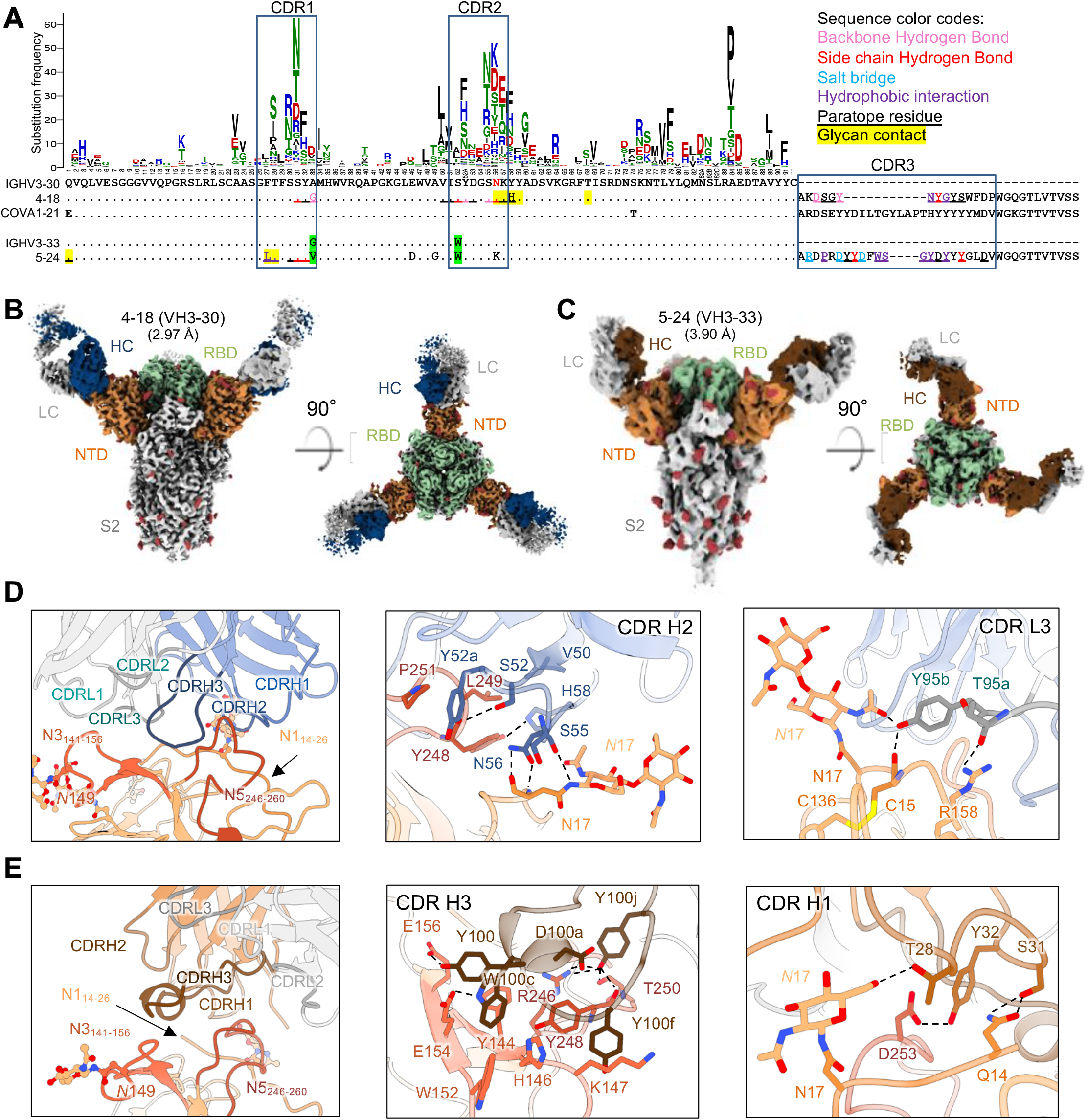
NTD-directed neutralizing antibodies derived from the closely related VH3-30 and VH3-33 genes show distinct binding modes. (A) Sequence alignment of VH3-30-derived (4-18) and VH3-33-derived (5-24) NTD-directed antibodies showing paratope residues, somatic hypermutations, and gene-specific substitution profile (GSSP) showing positional somatic hypermutation probabilities for VH3-30 gene. Substitutions between VH3-30 and VH3-33 germline genes are highlighted in green. (B) Cryo-EM reconstruction for spike complex with antibody 4-18 from two orthogonal views; NTD is shown in orange, RBD in green, glycans in red, with antibody heavy chain in blue and light chain in gray. (C) Cryo-EM reconstruction for spike complex with antibody 5-24 from two orthogonal views; NTD is shown in orange, RBD in green, glycans in red, with antibody heavy chain in brown and light chain in gray. (D) Expanded view of 4-18 interactions with NTD showing the overall interface (left panel), recognition in CDR H2 (middle panel), and recognition in CDR L3 (right panel). NTD regions N1 (residues 14-26), N3 (residues 141-156) and N5 (residues 246-260) are shown in shades of orange; CDR H1, H2, H3 are shown in shades of blue; CDR L1, L2, and L3 are shown in shades of gray. (E) Expanded view of 5-24 interactions with NTD showing the overall interface (left panel), recognition in CDR H3 (middle panel), and recognition in CDR H1 (right panel), colored as in (D) except for CDR H1, H2, H3, which are colored in shades of brown. See also Figure S1, Figure S3, Figure S6 and Table S1.

The cryo-EM structure of antibody 4-18 in complex with spike reveals an epitope that overlaps the VH1-24 antibodies, but with a significantly different overall mode of recognition. Overall, interactions are primarily mediated by CDR H2 and CDR L3, with additional contributions from CDRs H3, L1, and L2 (**Figure 2D, left panel; Figure S4B**). While CDR H3 inserts between NTD loops N3 and N5 like VH1-24 antibodies, the light chain CDR L3 binds adjacent to this region and also forms interactions with the NTD N1 loop. CDR H2 mediates extensive hydrogen bonding and hydrophobic interactions with NTD (**Figure 2D, middle panel**). CDR H2 residues Ser55_HC_ and Asn56_HC_ form hydrogen bonds with both backbone and side chain of Asn17_NTD_ at the base of the NTD *N*17 glycan in the N1 region. CDR H2 residues Ser52_HC_ and His58_HC_ also form hydrogen bonds with Tyr248_NTD_ in N5. Hydrophobic interactions are observed for CDR H2 residue Val50_HC_ with Leu249_NTD_ and Tyr52a_HC_ with Pro251_NTD_ in N5. In the light chain, Tyr95b_LC_ from the CDR L3 loop hydrogen bonds with Glycan *N*17 (**Figure 2D, right panel**) within the N-terminal region, which is typically disordered in ligand-free spikes.

Despite containing multiple aromatic residues in CDR H3, for the most part these residues do not form substantial hydrophobic interactions with residues from NTD, with exceptions Tyr98_HC_ and Tyr100_HC_, which bury 98 Å^2^ and 205 Å^2^ accessible surface area in the interface, respectively (**Figure S4B, left panel)**. Rather, the primary interactions mediated by CDR H3 are hydrogen bonds, including from the backbone carbonyl of Tyr98_HC_ with the backbone amino group of NTD Ser247_NTD_ and a hydrogen bond from the side chain hydroxyl of Tyr100_HC_ with the side chains from NTD residues Glu156_NTD_ and Arg158_NTD_.

The structure of VH3-33-derived antibody 5-24 (**Figure 2C**) in complex with spike reveals targeting of an overlapping epitope in NTD, but with overall recognition to be mediated by CDR H3 with additional contributions from CDR H1, but without the involvement of CDR L3 as seen for antibody 4-18 (**Figure 2E, left panel**). Also different from 4-18, four aromatic residues in the CDR H3 region of antibody 5-24 make extensive hydrophobic contacts with NTD loop N5 and the stem of the N3 β-hairpin (**Figure 2E, middle panel**), distinct from the hydrogen bond-dominated recognition observed in 4-18, with recognition by other CDRs also different.

Overall, while they target overlapping regions in NTD, recognition by VH3-33-deriveved antibody 5-24 is substantially different from that mediated by VH3-30-derived antibody 4-18. These dissimilarities show that, despite their derivation from highly similar VH genes, the NTD-directed neutralizing antibodies from VH3-30 and VH3-33 are not members of a single antibody class. Structural analysis showed that Ser52_HC_ from VH3-30 forms a hydrogen bond with Tyr248_NTD_. Substitution of Ser52_HC_ with the VH3-33-encoded Trp would lead to significant clashes with residues in CDR H2 and NTD loops N1 and N5 (**Figure S4C**), which could abolish the interaction between 4-18 and NTD. This suggests that the VH3-33 antibodies containing Trp52_HC_ cannot recognize NTD through a binding mode similar to VH3-30 antibody 4-18.

### NTD-directed neutralizing antibodies derived from the VH1-69 gene appear to comprise both reproducible and distinct classes

Of the 17 currently characterized NTD-directed or likely NTD-directed neutralizing antibodies (**Figure S1A**), three – antibodies 2-17 and 4-8 (Liu et al., 2020a) and antibody COV2-2676 (Zost et al., 2020) derived from the VH1-69 gene, with antibodies 2-17 and 4-8 deriving from the VH1-69*01 and VH1-69*02 alleles, respectively. Further, the CDR H3 regions of these antibodies showed similarity (**Figures 3A**). To understand the recognition of these VH1-69-derived antibodies we determined cryo-EM structures for spike complexes with the two most potent: antibodies 2-17 and 4-8 with IC_50_ potencies of 0.007 and 0.009 μg/mL.

**Figure 3.**
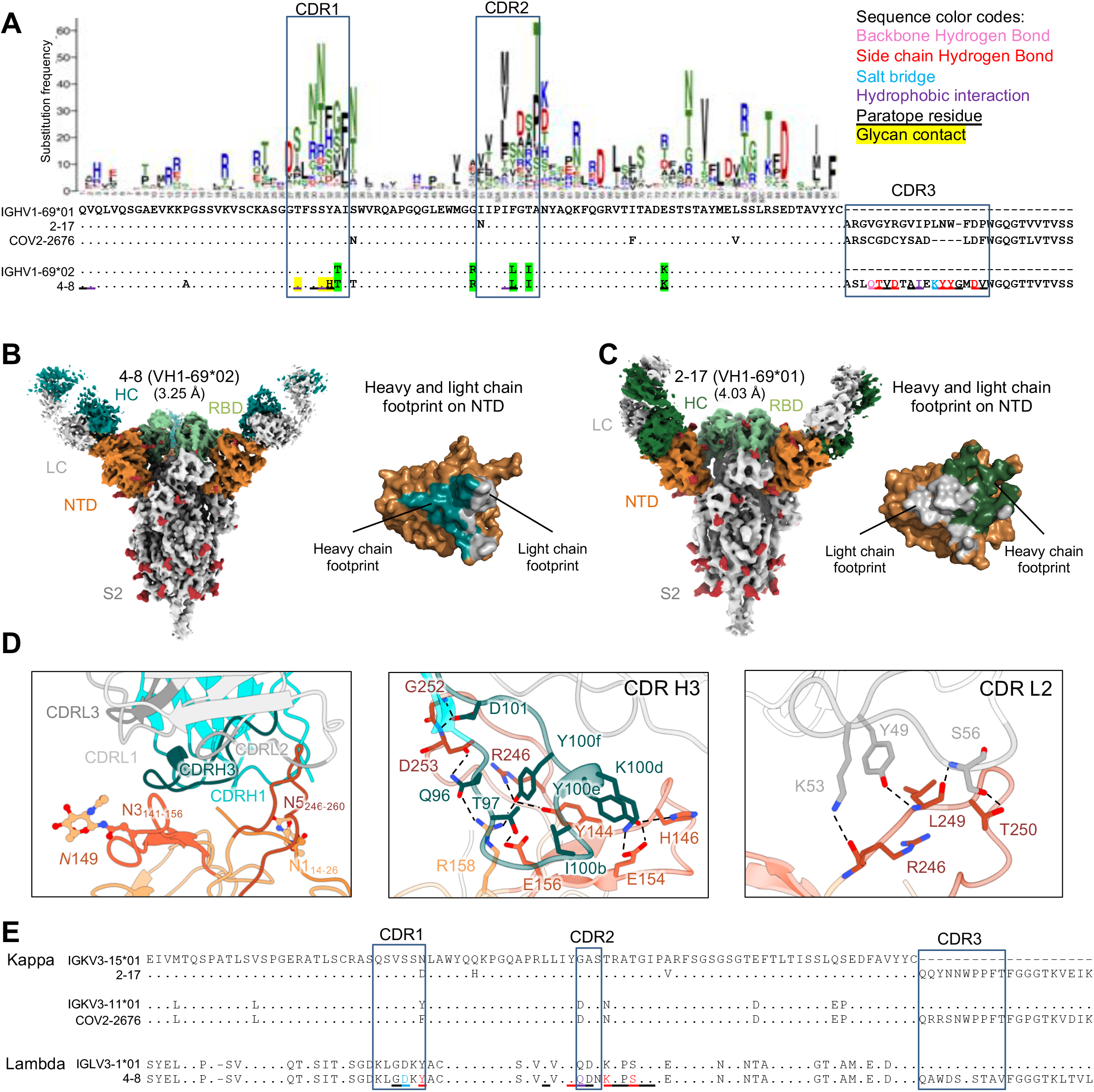
NTD-directed neutralizing antibodies derived from the closely related VH1-69*01 and VH1-69*02 genes show distinct binding modes. (A) Sequence alignment for VH1-69*01-derived (2-17) and VH1-69*02-derived (4-8) NTD-directed antibodies showing somatic hypermutations and paratope residues, with gene-specific substitution profile (GSSP) showing positional somatic hypermutation probabilities for VH1-69. Residues that differ between VH1-69*01 and VH1-69*02 alleles are highlighted in green. (B) Cryo-EM reconstruction for spike complex with antibody 4-8; NTD is shown in orange, RBD in green, glycans in red, with antibody heavy chain in teal and light chain in gray. Heavy and light chain footprint on NTD (right panel). (C) Cryo-EM reconstruction for spike complex with antibody 2-17 (left panel); NTD is shown in orange, RBD in green, glycans in red, with antibody heavy chain in dark green and light chain in gray. Heavy and light chain footprint on NTD (right panel) (D) Expanded view of 4-8 interactions with NTD showing the overall interface (left panel), recognition in CDR H3 (middle panel), and recognition in CDR L2 (right panel). NTD regions N1 (residues 14-26), N3 (residues 141-156) and N5 (residues 246-260) are shown in shades of orange; CDR H1, H2, H3 are shown in shades of teal; CDR L1, L2, and L3 are shown in shades of gray. (E) Sequence alignment of light chain of VH1-69-derived antibodies showing diverse germline gene usage (2-17 and COV2-2676 utilizes kappa light chain, 4-8 utilizes lambda light chain), IGKV3-15*01 is used as reference. Paratope residues of 4-8 arecolored as in (A). See also Figure S1, Figure S3, Figure S6 and Table S1.

Single-particle cryo-EM data for antibody 4-8 yielded a 3D reconstruction at 3.25 Å resolution (**Figure 3B, Figure S3D and Table S1**), however, like antibody 2-17 (**Figure 3C, Table S1**), and as reported previously (Liu et al., 2020a), high mobility of the bound Fab blurred the interface region. We used local refinement with particle subtraction to obtain a high-quality reconstruction for the 4-8 interface with spike (**Figure 3D**). Like the VH1-24-derived antibodies, CDR H3 binds between the NTD N3 and N5 loops, but in a distinctive way; CDR H3 dominates the interface and its approach to the N3/N5 region is nearly orthogonal to that observed for CDR H3 in the VH1-24 class antibodies (**Figure 3D, middle panel**). Recognition by other CDRs is also distinct from the other NTD-directed neutralizing antibodies (**Figure 3D, right panel; Figure S4D**)

We also collected single-particle cryo-EM data for antibody 2-17, yielding a 3D reconstruction at 4 Å resolution (**Figure 3C, Table S1**). Despite the poor resolution of the 2-17 cryo-EM maps, particularly in the interface region, we were able to dock a threaded 2-17 structure into the cryo-EM reconstruction of the complex with high confidence.

We compared the heavy and light chain epitope footprints on NTD for antibodies 4-18 (**Figure 3B, right panel**) and 2-17 (**Figure 3C, right panel**). Notably, the orientation of heavy and light chains between 2-17 and 4-8 were rotated ∼90 degrees from each other, indicating different modes of recognition, and showing that they are of different classes.

We next asked whether other NTD-directed neutralizing antibodies derived from VH1-69 heavy chain might be members of the 2-17 or 4-8 classes. Analysis of light chains indicated antibodies 2-17 and COV2-2676, derived from genes KV3-15 and KV3-11, respectively, to be remarkably similar in their light chain CDR L3 regions suggesting that 2-17 and COV2-2676 might be of the same class. Overall, we observed NTD-directed neutralizing antibodies from VH1-69 to form at least two classes, with similarity of light chains suggesting antibodies 2-17 and COV2-2676 might represent a reproducible class observed in two different donors.

### Functional requirements for somatic hypermutation (SHM)

Sequence analyses showed that all NTD-directed antibodies accumulate somatic hypermutations in their paratope regions (**Figures 1A, 2A, and 3A**). To help understand characteristics of the antibody precursors, we reverted the paratope-region somatic hypermutations observed in seven NTD-directed antibodies (1-87, 1-68, 2-51, 4-18, 5-24, 4-8, and 2-17) to their respective germline residues. Overall, these germline-reverted antibodies showed significantly reduced binding affinities and neutralization potencies (**Figure S6**). For the VH1-24 multi-donor antibody class, antibodies 1-87, 1-68, and 2-51 shared two convergent somatic hypermutations (T30I and G55A/V) in heavy chain, reversion of which showed significantly (∼4-20-fold) reduced binding affinity (**Figure S6A-B**). For antibodies 4-18 and 5-24, derived from VH3-30 and VH3-33, respectively, reversion of SHMs in combination in each antibody nearly abolished neutralization (**Figure S6C**). For the two VH1-69-derived antibodies 2-17 and 4-8, neutralization was improved by SHMs by ∼20- and ∼8-fold, respectively. Thus, all of the NTD-directed potently neutralizing antibodies we tested required affinity maturation to achieve high binding affinity and high potency. The VH3-30 and VH3-33 antibodies were more sensitive to SHMs than the antibodies derived from other VH genes. Nonetheless, antibodies corresponding to the initial recombinants, with reversion of all paratope SHMs in combination, could still bind to spike with apparent IgG K_D_s ∼2-70 nM, suggesting that precursor B cells of the NTD antibodies are likely to be efficiently activated by spike binding. Gene-specific substitution profiles (Sheng et al., 2017) showed that the observed SHMs are each generated by the somatic hypermutation machinery with high frequencies (**Figures 1A, 2A, and 3A**), suggesting that requirements for SHM are unlikely to present a significant barrier to antibody development.

### NTD-directed potently neutralizing antibodies have similar angles of approach

To gain an overall understanding of the angle of antibody approach to the spike by these NTD-directed potently neutralizing antibodies, we determined their angles of approach around a latitudinal axis to define freedom between viral and host cell membranes, and around a longitudinal axis to define freedom within the plane of the membrane. Relative to the viral spike, the latitudinal axis is perpendicular to the trimer axis, and the longitudinal axis is parallel to this axis. Latitudinal and longitudinal approach angles among the NTD-neutralizing antibodies were similar – with antibodies approaching spike with antigen-combining surface oriented towards the viral membrane (**Figure 4A-C**).

**Figure 4.**
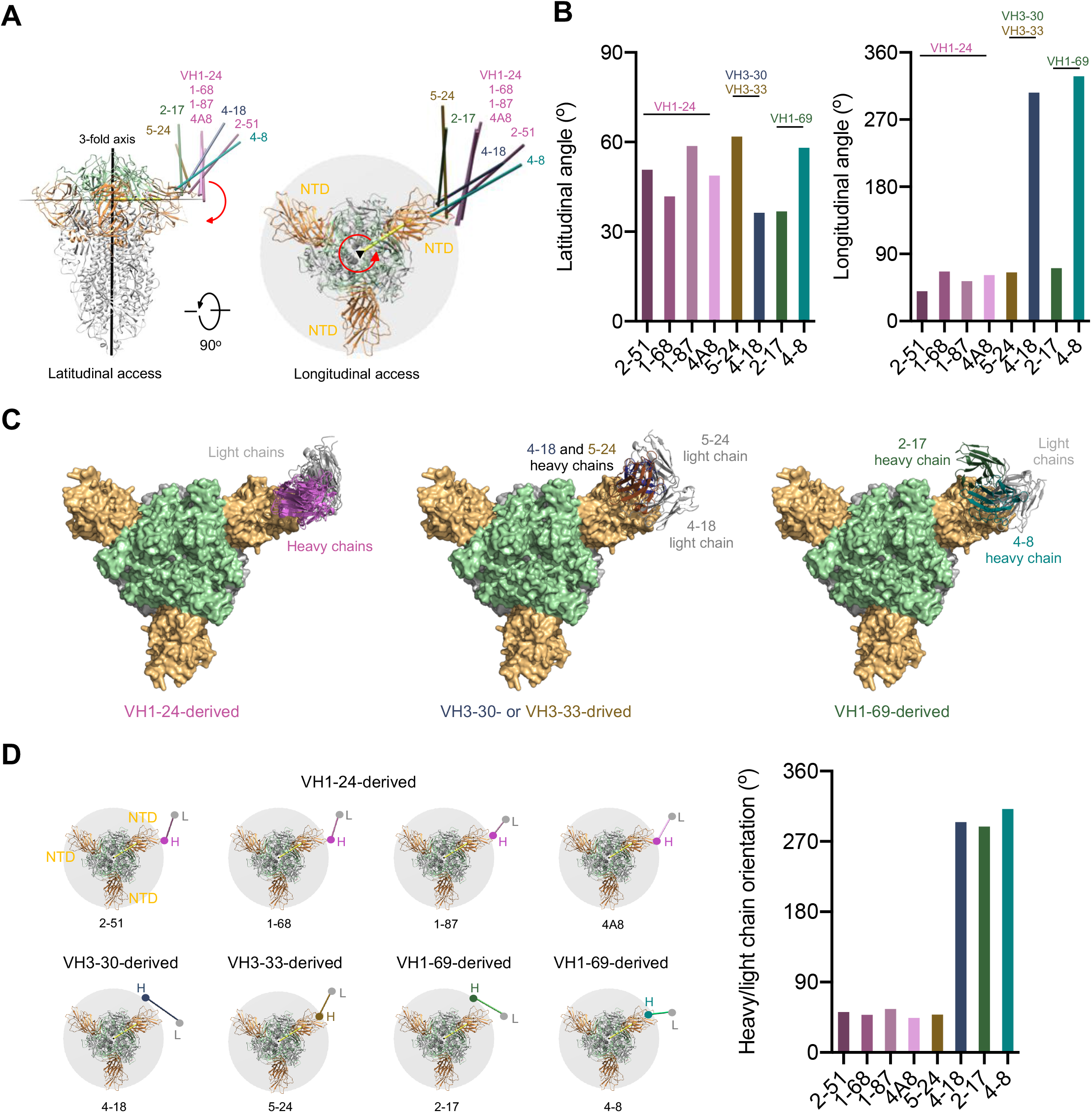
Angles of approach for NTD-directed neutralizing antibodies. (A) Overall approach of NTD-directed neutralizing antibodies to spike with angles defined with red arrows. Three-fold axis is indicated by a black triangle. Antibodies are represented by long axes of the Fabs and colored by heavy chain colors defined in Figures 1-3. (B) Latitudinal and longitudinal angles of approach. (C) Angles of recognition for antibodies grouped by VH-gene. Notably, only those from VH1-24 show a consistent orientation. (D) Heavy-light chain orientations show graphically (left) and quantitatively (right).

We also analyzed the heavy-light chain orientations in the complexes with spike (**Figure 4D**). Here the NTD-directed antibodies differed, with three of the antibodies, 4-18 from VH3-30 and 2-17 and 4-8 from VH1-69, showing heavy and light chain angles of approach that differed from the other five antibodies. Thus, while the heavy/light orientation relative to spike could differ substantially, lesser differences were observed in latitudinal and longitudinal angles of approach, with all NTD-neutralizing antibodies approaching spike from “above” with their antigen-binding surfaces oriented towards the viral membrane.

### NTD-directed antibodies induce conformational changes in NTD and spike

To gain insight into the impact of antibody recognition on the conformation of NTD, we superimposed antibody-spike or antibody-NTD complexes onto the NTD domain, and examined the structural alteration in NTD versus NTD in the ligand-free spike (PDB: 6ZGE), calculating the per-residue Cα movement between bound and ligand-free (**Figure 5A**), which ranged as high as 16-18 Å for most of the NTD-directed antibodies, though 4-8 (10.1 Å) and 5-24 (11.7 Å) were somewhat lower. The largest structural change occurred in the N3 β-hairpin, although the mobile N1 and N5 loops also showed large deviations (**Figure 5B**). In general, the regions of NTD that moved were contacted by antibody (**Figure 5C**), indicating the conformational changes were a direct consequence of antibody binding. In addition to the conformational change induced in NTD, we observed other changes in spike. Notably, the 4-18 antibody bound spike was substantially better ordered than the other NTD-bound spikes (achieving a nominal cryo-EM resolution of 2.97 Å, which was ∼1 Å better than most of the other complexes). Examination of the 4-18 bound spike indicated almost 40° rotation in the central S2 triple helical bundle (**Figure S4E**).

**Figure 5.**
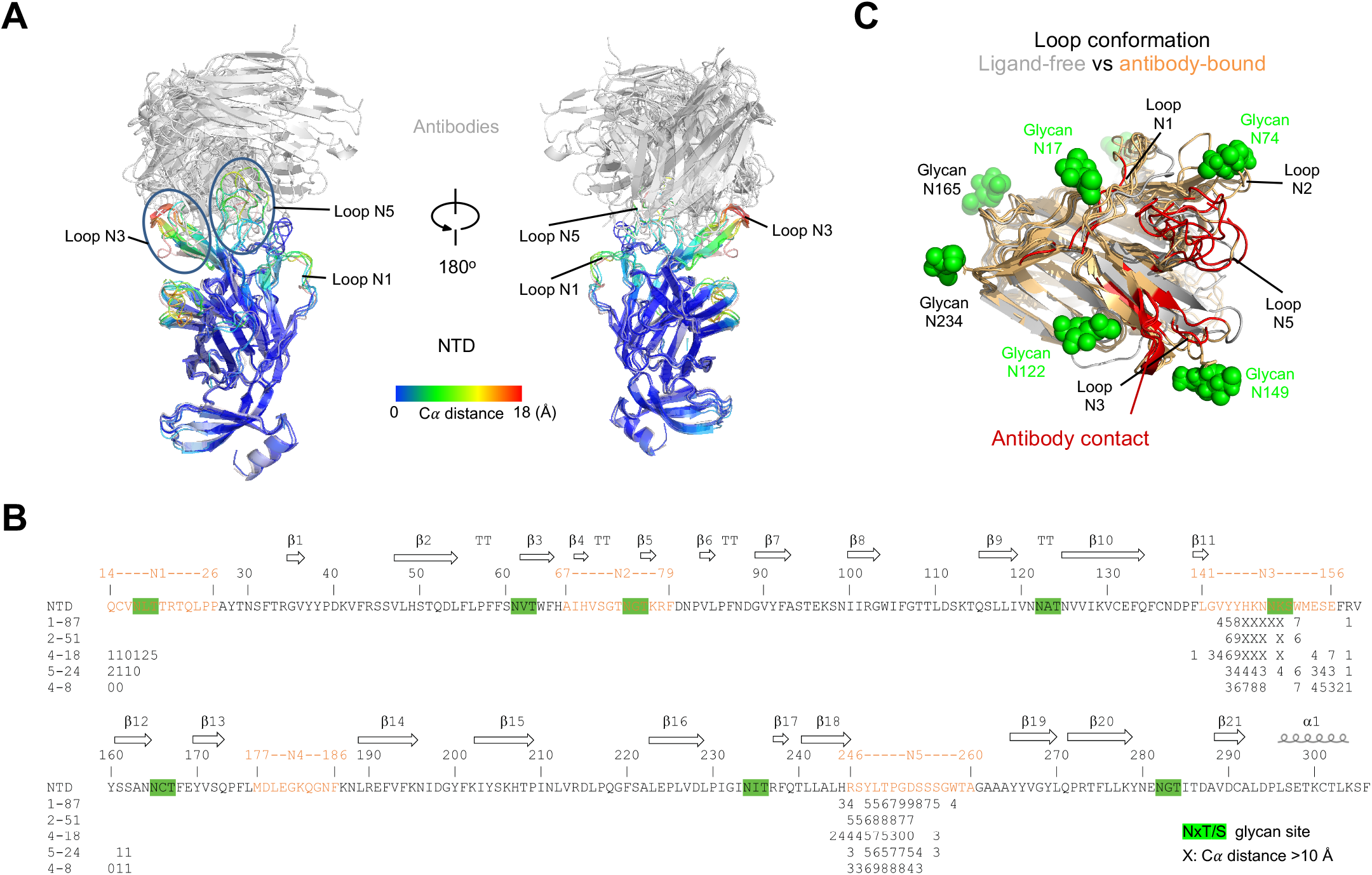
NTD-directed antibodies induce conformational changes in NTD and spike. (A) Conformational changes in NTD induced by binding of neutralizing antibodies. Antibody-bound NTDs are shown in cartoon representation and colored by per residue Cα movements compared to unliganded NTD. Antibodies are shown in gray cartoon. Major NTD loops interacting with antibodies are labeled. (B) Sequence of NTD highlighting antibody contact and conformational change. Epitope residues for each antibody are marked with a number representing C*α* movements (Å) from unliganded NTD, the symbol “X” indicates movement 10 Å and above. Potential glycosylation sites on NTD are highlighted in green. (C) Epitope regions on NTD (red) and their conformational change. Glycans on NTD are shown as green spheres.

Overall, binding of NTD-directed antibodies induced substantial structural rearrangements, not only in recognized loops but also of the N3-β-hairpin. The higher immunogenicity observed with flexible regions likely stems from the ability of these regions to assume distinct conformations required for diverse antibodies to bind, with the recognized site on NTD apparently exemplifying this effect.

### The NTD supersite

To define the spike surface recognized by potent NTD-directed neutralizing antibodies, we analyzed the epitopes for all eight of the NTD-directed neutralizing antibodies with defined structures, the seven described in this study as well as antibody 4A8, described previously (Chi et al., 2020) These ranged in potency from remarkably potent 2-17 and 2-51 antibodies with IC_50_ of 0.007 μg/mL to the neutralizing, but substantially less potent 4A8 with IC_50_ of 0.39 μg/mL; all eight of these antibodies recognize overlapping epitopes on NTD (**Figure 6A**). We define the spike surface recognized by at least two antibodies, from different classes, of these eight potent NTD-directed antibodies, as the NTD supersite (**Figure 6B**).

**Figure 6.**
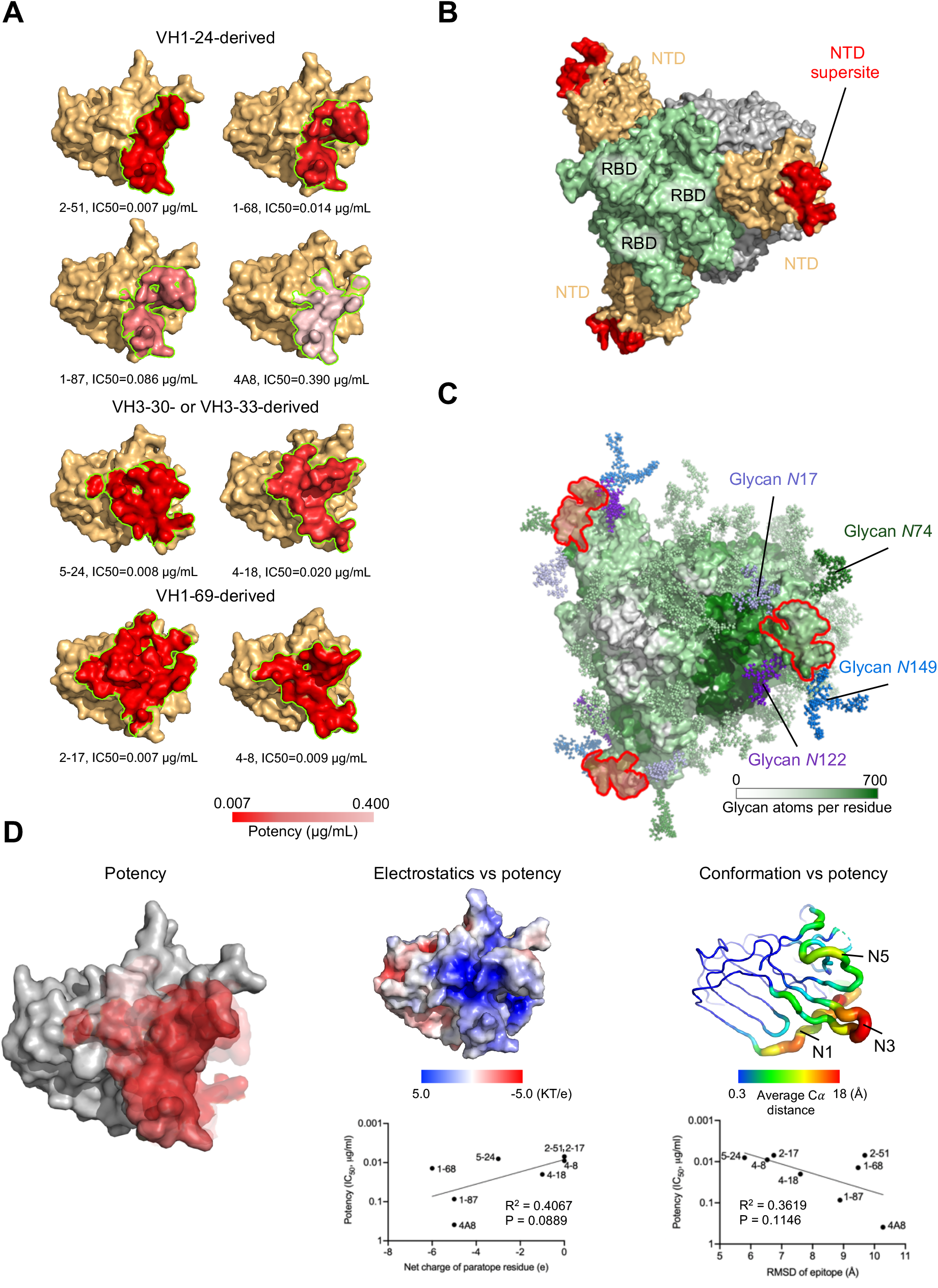
A structurally plastic antigenic supersite in the distal-loop region of NTD revealed by comparison of antibodies derived from the four multi-donor classes. (A) Epitopes of NTD-targeting antibodies colored by potency. (B) The supersite of vulnerability on NTD. (C) Glycan coverage of the spike. The NTD supersite is surrounded by glycans at *N*17, *N*74, *N*122 and *N*149. (D) NTD structural properties and antibody potency. Epitope surfaces of different antibodies were overlaid onto NTD with shades of red representing potency (left). Trending correlations were identified between antibody potency and epitope electrostatics (middle) and conformational change (right). See also Figure S1, Figure S6, Figure S7 and Table S3.

The NTD supersite was located at the periphery of the spike, distal from the 3-fold axis, and facing away from the viral membrane. This surface was surrounded by four glycans, *N*17, *N*74, *N*122, and *N*149, and nominally ‘glycan free’, although molecular dynamics simulations with fully-glycosylated spike indicated some glycan-coverage, though less than adjacent regions more proximal to the spike 3-fold (**Figure 6C**).

To gain insight into the structural features of the NTD supersite, we first analyzed the distribution of epitopes versus potency, but did not observe substantial variation in potency over the NTD supersite (**Figure 6D, left panel)**. In addition, we found no correlation between Fab affinity (**Table S3**) and potency. Electrostatic surface analysis revealed the supersite to have strong positive electrostatic potential (**Figure 6D, middle panel**), while recognizing antibodies had complementary strong electronegative potentials (**Figure S7**). Interestingly, we observed the potency of the antibody to trend with decreasing net negative charge on the paratope, which was surprising as complementary electrostatics would be expected to enhance affinity, and thus potency.

With respect to recognized conformation, we compared the ligand-free conformation of the supersite versus its antibody-bound conformation; the recognized β-strands at the center of the epitope were displaced ∼3 Å, with loops N1, N3 and N5 moving substantially more, up to 18 Å (**Figure 6D, right panel**). With respect to correlation with potency, we observed the magnitude of induced NTD conformational change to trend inversely with potency. This is not surprising in that the requirement for conformational change is likely to lower the energy of binding. Notably, antibodies that induced larger conformational changes were also more electronegative, potentially providing an explanation for the observation that increasing negative charge trended with reduced potency.

Overall, the NTD supersite comprised a structurally plastic surface, formed primarily by the N3 β-hairpin and including other flexible regions such as the N1 and N5 loops. This surface was also both glycan-free and highly electropositive – and facing away from the electronegative viral membrane.

### NTD supersites in other betacoronaviruses

To understand the generality of the single-NTD supersite that we observe for SARS-CoV-2, we examined the recognition of NTD-directed neutralizing antibodies targeting other betacoronaviruses. Searches of the PDB found only two NTD-directed antibodies targeting other betacoronaviruses: these two antibodies, G2 (Wang et al., 2018) and 7d10 (Zhou et al., 2019), both neutralized MERS and targeted overlapping glycan-free surfaces on NTD facing away from the viral membrane (**Figure 7A**). The epitopes for both of these antibodies partially overlapped the analogous surface comprising the NTD-supersite in SARS-CoV-2, but were more centrally located and more proximal to the spike 3-fold.

**Figure 7.**
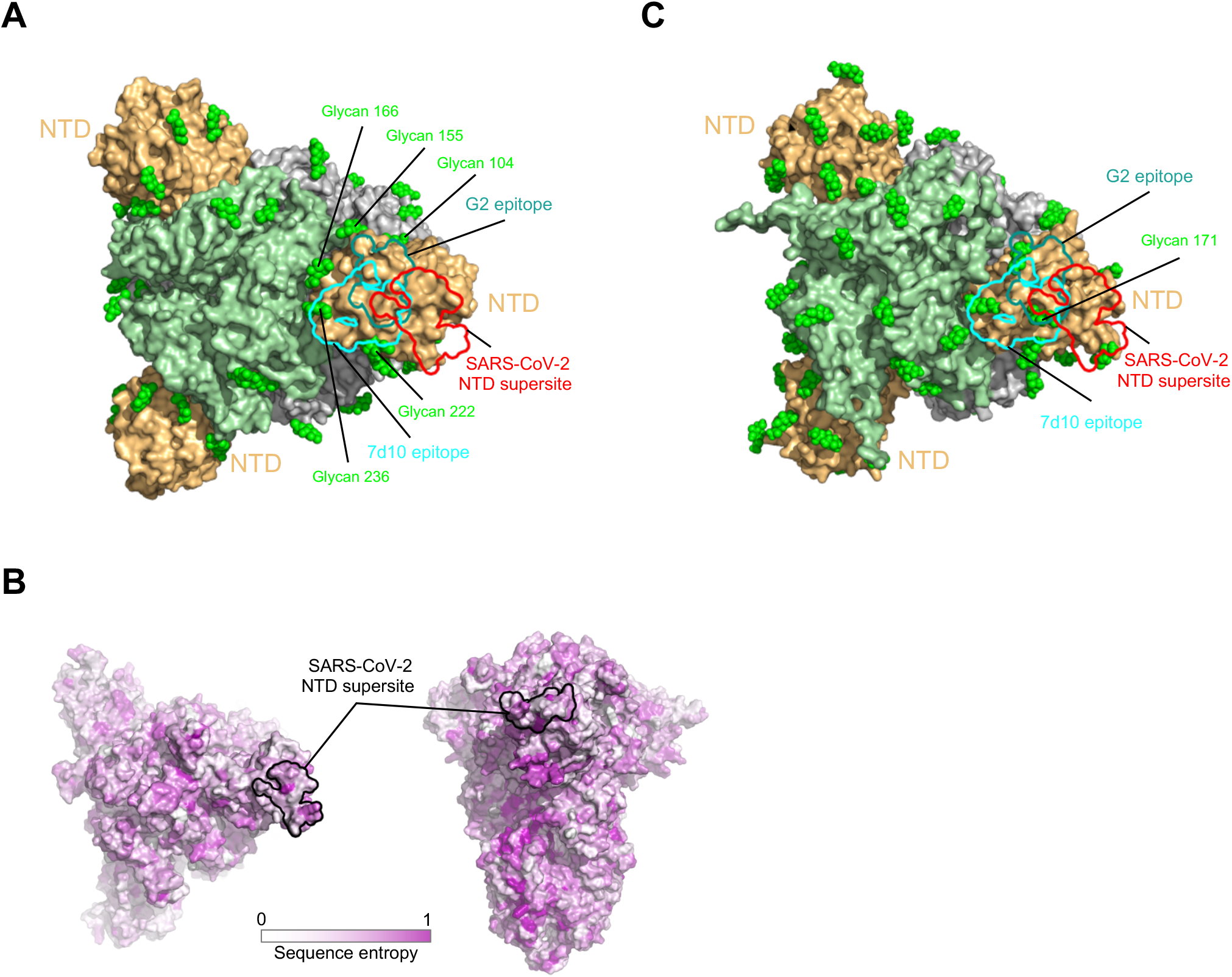
NTD supersite on MERS betacoronaviruses. (A) Epitopes of MERS NTD antibodies target a site closer to the trimer axis. Borders of epitopes of antibody G2 and 7d10 are colored teal and cyan, respectively. SARS-CoV-2 NTD supersite is show as red boundary line. Glycans are shown as green spheres. (B) Spike sequence entropy between betacoronaviruses. (C) NTD of HKU1 spike is substantially glycosylated.

Quantification of the average number of proximal glycan atoms indicated lower glycan density over the NTD-supersite than over the equivalent surfaces recognized by G2 and 7d10 antibodies. The epitopes recognized by these two antibodies also showed substantially less conformational mobility. Thus, potent NTD-directed neutralizing antibodies targeting SARS-CoV-2 preferentially recognized a less glycosylated, more flexible region than the analogous surfaces recognized by NTD-directed antibodies neutralizing MERS.

Since the MERS and SARS-CoV-2 directed antibodies both targeted glycan-free sites on NTD facing away from the viral membrane, we sought to understand the properties of the equivalent surfaces of other coronavirus spikes. We calculated the sequence divergence of other betacoronoviruses and mapped this to the NTD surface, which showed high diversity in sequence (**Figure 7B**). We modeled the sequence-predicted glycans on the HKU1 spike structure (PDB: 5I08) and examined the location of glycans on HKU1 NTD (**Figure 7C**). Notably glycan *N*171 were observed to be directly the region of overlap between the equivalent positions of the SARS-CoV-2 supersite, 7d10 and G2 epitopes. Thus, the presence of glycans may impact the presence of absence of NTD-sites of vulnerability in betacoronaviruses.

## Discussion

Antibodies directed to NTD and to RBD can neutralize with high potency (less than 0.01µ g/mL IC_50_). While RBD shows many non-overlapping sites of vulnerability to antibody (Barnes et al., 2020a; Brouwer et al., 2020; Lv et al., 2020; Pinto et al., 2020b; Yuan et al., 2020a), NTD appears to contain only a single site of vulnerability to neutralization. As discussed above, one reason for this may be the high glycan-density on NTD, with 8 *N*-linked glycans in ∼300 residues, a density of one glycan per ∼40 residues, and few glycan-free surfaces that can be easily recognized by the immune system. A second reason may be the restricted approach angle which we observed for all known NTD-directed neutralizing antibodies, including the seven reported here, which all approach spike from “above”. We note in this context that competition analysis indicates other NTD-directed antibodies capable of recognizing spike and forming a separate competition group to be non-neutralizing (Liu et al., 2020) – and the other large surface on NTD that is exposed on spike faces toward the viral membrane. This surface is mostly glycan-free, and antibodies binding to it would be required to approach from “below”. Thus, unlike RBD, where neutralizing antibodies appear to have diverse approach angles, the presence of only a single-NTD site of vulnerability may relate to the requirement to approach from “above.”

In addition to satisfying requirements stemming from the restricted approach angle, the higher relative prevalence of NTD-supersite-directed antibodies is likely to stem from increased immunogenicity due to both the lower relative glycan density of the supersite, as well as the flexible nature of the N3 hairpin and N5 loop primary recognition regions, and their ability to assume distinct conformations that allow for recognition by diverse antibodies. In the case of the multi-donor VH1-24 antibody class, which arises from the most prevalent VH-gene utilized (**Figure S1A**), we found VH1-24 to be the most negatively charged human VH gene **(Figure S5B)**. Such negative electrostatic potential complements the highly electropositive nature of the NTD supersite that we observe here (**Figure 6D**). Thus, multiple factors including epitope glycosylation and flexibility, restrictions on approach angle, and paratope charge complementarity can contribute to the prevalence of antibodies targeting the NTD supersite.

Although the approach to the spike from “above” observed for all NTD-directed neutralizing antibodies is consistent with a neutralization mechanism based on steric hindrance of spike interaction with ACE2 receptor at the cell membrane, there is currently no evidence for competition between NTD-directed antibodies and ACE2 (Liu et al., 2020a). A plausible alternative model would be for antibody recognition of the NTD supersite to impede spike function in mediating fusion of virus and host cell membranes. Indeed, protease-resistance analysis of MERS spike in complex with MERS NTD-directed neutralizing antibody 7d10 showed that 7d10 binding prevented increased protease sensitivity associated with the prefusion-to-postfusion transition (Zhou et al., 2019); further studies will be required to understand mechanisms of neutralization for antibodies that recognize the NTD-supersite.

With respect to vaccine implications, our results clearly identify the NTD site of vulnerability most likely to elicit neutralizing antibodies. There are many ways that this information can be incorporated into vaccine design, including the inclusion of NTD along with RBD in vaccine formulations, the multivalent display of the NTD supersite on nanoparticle immunogens, and epitope-focusing through the creation of scaffolds displaying the N3-β-hairpin and other regions of recognized by NTD-directed neutralizing antibodies.

With respect to the therapeutic potential of NTD-directed antibodies, these target a site that is remote from those targeting RBD sites and thus should provide complementary neutralization to RBD-directed antibodies and require distinct escape pathways. The fact that all, or a great majority, of NTD-neutralizing antibodies target a single site, however, suggests there may be little utility to utilizing combinations of NTD-directed neutralizing antibodies.

## Acknowledgments

We thank R. Grassucci, Y.-C. Chi and Z. Zhang from the Cryo-EM Center at Columbia University for assistance with cryo-EM data collection, R. Amaro and L. Casalino for providing molecular dynamics trajectories before publication, P. Tripathi for provision of 4A8 and 2-17 Fabs, B. Zhang for provision of 4A8, T. Bylund for help with bioinformatics and members of the Virology Laboratory and Vector Core, Vaccine Research Center, for discussions and comments on the manuscript. We thank D. Neau, S. Banerjee, and S. Narayanasami for help with synchrotron data collection conducted at the APS NE-CAT 24-ID-C beamline, which is supported by National Institutes of Health (NIH) P41 GM103403; use of NE-CAT at the Advanced Photon Source was supported by the US Department of Energy, Basic Energy Sciences, Office of Science, under contract number W-31-109-Eng-38. Support for this work was provided by the Intramural Research Program of the Vaccine Research Center, National Institute of Allergy and Infectious Diseases (NIAID). Support for this work was also provided by Samuel Yin, Pony Ma, Peggy & Andrew Cherng, Brii Bioscieces, Jack Ma Foundation, JBP Foundation, Carol Ludwig, and Roger & David Wu, COVID-19 Fast Grants, the Self Graduate Fellowship Program, and NIH grants DP5OD023118, R21AI143407, and R21AI144408. Some of this work was performed at the Columbia University Cryo-EM Center at the Zuckerman Institute, and some at the Simons Electron Microscopy Center (SEMC) and National Center for Cryo-EM Access and Training (NCCAT) located at the New York Structural Biology Center, supported by grants from the Simons Foundation (SF349247), NYSTAR, and the NIH National Institute of General Medical Sciences (GM103310).

## Author Contributions

GC led the determination of 5 cryo-EM structures shown in Fig. 1-3, YG provided sequence analyses defining classes in Fig. 1-3, TZ led analysis for Fig. 4-7, JG carried out map refinement and model building for 4-8 and 2-17 structures with spike, ML provided glycan-masking analysis for supersite, correlations for supersite properties and threading of diverse spikes, MR led determination of 2 cryo-EM structures, ER led crystal structure determination of 2-51 with isolated NTD, JY provided antibodies for structural analysis, FB provided spike, mutant spikes and stabilized spikes, JB provided almost all Fabs, YH, LL, MSN and PW carried out antibody reversion analyses, PSK determined SPR affinity measurements, RR assisted with glycan density calculations, ASO provided spike, REA provided MSD trajectories for glycosylated spike, GYC supervised informatics analysis and assisted with antibody structural threading and docking, DDH supervised IgG production and reversion analysis, ZS supervised sequence analysis, PDK supervised structural analyses, and LS supervised structural determinations and led the overall project.

## Declaration of Interests

DDH, YH, JY, LL, MSN and PW are inventors of a patent describing some of the antibodies reported on here.

## RESOURCE AVAILABILITY

### Lead Contact

Further information and requests for resources and reagents should be directed to and will be fulfilled by Lawrence Shapiro.

### Materials Availability

Expression plasmids generated in this study for expressing SARS-CoV-2 proteins and antibody mutants will be shared upon request.

### Data and Code Availability

The cryo-EM structures and the crystallographic structure are in the process of being deposited to the Electron Microscopy Data Bank (EMDB) and the Protein Data Bank (RCSB PDB). Cryo-EM structural models and maps of NTD-directed antibodies in complex with SARS-CoV-2 spike have been deposited in the PDB and EMDB for antibodies 1-87 (PDB:7L2D, EMDB: EMD-23125), 4-18 (PDB:7L2E, EMDB: EMD-23126) and 5-24 (PDB: 7L2F, EMDB: EMD-23127); cryo-EM maps have been deposited for antibodies 1-68 (EMDB: EMD-23150) and 2-51 (EMDB: EMD-231251). The crystallographic structure of antibody 2-51 in complex with SARS-CoV-2 spike NTD has been deposited in the PDB with accession code 7L2C.

## EXPERIMENTAL MODEL AND SUBJECT DETAILS

### Cell lines

HEK 293T/17(cat# CRL-11268™), I1 mouse hybridoma (cat# CRL-2700) and Vero E6 cells (cat# CRL-1586™) were from ATCC. Expi293F™ Cells (cat# A39240) and Expi293F™ GnTI-Cells (cat# A39240) were from Thermo Fisher Scientific.

## METHOD DETAILS

### Protein Samples Expression and Purification

The SARS-CoV-2 S2P and HexaPro spike variant constructs were produced as described in Wrapp et al., 2020 and in Hsieh et al., 2020 respectively. They were expressed in Human Embryonic Kidney (HEK) 293 Freestyle cells (Invitrogen) in suspension culture using serum-free media (Invitrogen) and transfected into HEK293 cells using polyethyleneimine (Polysciences). Cell growths were harvested four days after transfection, and the secreted proteins were purified from supernatant by nickel affinity chromatography using Ni-NTA IMAC Sepharose 6 Fast Flow resin (GE Healthcare) followed by size exclusion chromatography on a Superdex 200 column (GE Healthcare) in 10 mM Tris, 150 mM NaCl, pH 7.4.

The N-terminal domain of SARS-CoV-2 spike (NTD, residues 1-330) was cloned into the pVRC-8400 mammalian expression plasmid, with a C-terminal 6X-His-tag cleavable by HRV-3C protease. The NTD construct was transiently transfected into HEK293 GnTI-Freestyle cells suspension culture in serum-free media using polyethyleneimine. Four days after transfection, the secreted protein was purified using Ni-NTA IMAC Sepharose 6 Fast Flow resin followed by size exclusion chromatography on a Superdex 200 column in 10 mM Tris, 150 mM NaCl, pH 7.4. Fractions containing NTD were combined and 1% (w/w) HRV-3C protease (Thermo fisher) was added to remove the C-terminal His-tag, followed by incubation for 24 hrs at 4 °C. Inverse IMAC using Ni-NTA resin was then performed to purify NTD from the His-tag and residual uncleaved protein. Enzymatic deglycosylation of NTD was carried out by adding 2.5 μL Endo Hf (NEB) per 20 μg of NTD and incubating for 24 hrs at 25 °C; a second round of SEC was performed to remove excess Endo Hf and to exchange buffer in 10 mM Tris, 150 mM NaCl, pH 7.4. Protein purity was analyzed by SDS-PAGE at every step.

NTD-directed monoclonal antibodies 1-68, 1-87, 2-17, 2-51, 4-8, 4-18 and 5-24 were expressed and purified as described in (Liu et al., 2020a).Fabs fragments were produced by digestion of IgGs with immobilized papain at 37 °C for 3 hrs in 50 mM phosphate buffer, 120 mM NaCl, 30 mM cysteine, 1 mM EDTA, pH 7. The resulting Fabs were either purified from Fc by affinity chromatography on protein A (1-68, 1-87, 2-17, 2-51 and 4-8) or used as Fab/Fc mixture (4-18 and 5-24). Fab purity was analyzed by SDS-PAGE; all Fabs were buffer-exchanged into 10 mM Tris, 150 mM, pH 7.4 for crystallization and cryo-EM experiments.

### Antibody mutagenesis

For each antibody, variable genes were optimized for human cell expression and synthesized by GenScript. VH and VL were inserted separately into plasmids (gWiz or pcDNA3.4) that encoding the constant region for heavy chain and light chain. Monoclonal antibodies were expressed in Expi293 (ThermoFisher, A14527) by co-transfection heavy chain and light chain expressing plasmids using polyethylenimine (PEI, Linear, MV∼25,000, Polysciences, Inc. Cat. No. 23966) and culture in 37 °C degree shaker at 125RPM and 8% CO2.

Supernatants were collected on day 5, antibodies were purified by rProtein A Sepharose (GE, 17-1279-01) affinity chromatography.

Antibody gene mutations were introduced by QuikChange II site directed mutagenesis kit (Agilent, Cat. No. 200524)

### Antibody Fab binding affinity measurement by Surface Plasmon Resonance

SPR binding assays for Fabs were performed using a Biacore T200 biosensor, equipped with a Series S CM5 chip, in a running buffer of 10 mM HEPES pH 7.4, 150 mM NaCl, 0.1 mg/mL BSA and 0.01% (v/v) Tween-20 at 25 °C.

HexaPro Spike was captured through its C-terminal his-tag over an anti-his antibody surface. These surfaces were generated using the His-capture kit (Cytiva, MA) according to the instructions of the manufacturer, resulting in approximately 10,000 RU of anti-his antibody over each surface. HexaPro was captured over a single flow cell at a capture level of 500-800RU with Fabs with higher KDs (2-17 and 4-18) requiring higher capture levels. An anti-his antibody surface was used as a reference flow cell to remove bulk shift changes from the binding signal. Fabs were tested using a three-fold dilution series ranging from 2.96-240 nM, except for Fabs 4-18 and 2-17, which were analyzed at concentrations of 8.88-720 nM. The association and dissociation rates were each monitored for 120 s and 600 s respectively, at 50 μL/min. The bound HexaPro/Fab complex was regenerated from the anti-his antibody surface using a 10s pulse of 15 mM H_3_PO_4_ at a flow rate of 100 μL/min, followed by a 60 s buffer wash at the same flow rate. Each Fab was tested in order of increasing protein concentration, in duplicate. Blank buffer cycles were performed by injecting running buffer instead of Fab to remove systematic noise from the binding signal. The data was processed and fit to 1:1 single cycle model using the Scrubber 2.0 (BioLogic Software).

### Full IgG binding Affinity Measurements by Surface Plasmon Resonance

The mammalian expression vector that encodes the ectodomain of the SARS-CoV-2 S trimer for full IgG binding affinity measurement was kindly provided by Dr. Jason McLellan (Wrapp et al., 2020b). SARS-CoV-2 S trimer expression vector was transiently transfected into Expi293TM cells using 1 mg/mL of polyethylenimine (Polysciences). Five days post transfection, the S trimer was purified using Strep-Tactin XT Resin (Zymo Research).

The binding affinities of full IgG antibodies to SARS-CoV-2 spike protein were determined using surface plasmon resonance (SPR) and a BIAcore T200 instrument (GE Healthcare) at 25°C. The anti-his antibody was first immobilized onto two different flow cells of a CM5 sensorchip (BR100030, Cyvita) surface using the His Capture Kit (28995056, Cyvita) according to the manufacturer’s protocol. The His-tagged SARS-CoV-2 spike protein was then injected and captured on flow cells 2. Flow cells 1 was used as the negative control. A three-fold dilution series of antibodies with concentrations ranging from 300 nM to 1.2 nM were injected over the sensor surface for 30 s at a flow rate of 10 µL/minute. The dissociation was monitored for 300 s and the surface was regenerated with 10 mM Glycine pH 1.5 (BR100354, Cyvita). The running and sample buffer is 10 mM HEPES pH 7.4, 150 mM NaCl, 3 mM EDTA, 0.05% P-20 (HBS-EP+ buffer, BR100826, Cyvita). The resulting data were fit to a 1:1 binding model using Biacore Evaluation Software and were plotted using Graphpad.

### Neutralization assays

Recombinant Indiana VSV (rVSV) expressing SARS-CoV-2 spikes were generated as previously described. HEK293T cells were grown to 80% confluency before transfection with pCMV3-SARS-CoV-2-spike (kindly provided by Dr. Peihui Wang, Shandong University, China) using FuGENE 6 (Promega). Cells were cultured overnight at 37 °C with 5% CO_2_. The next day, medium was removed and VSV-G pseudo-typed ΔG-luciferase (G*ΔG-luciferase, Kerafast) was used to infect the cells in DMEM at a MOI of 3 for 1 hr before washing the cells with 1X DPBS three times. DMEM supplemented with anti-VSV-G antibody (I1, mouse hybridoma supernatant from CRL-2700; ATCC) was added to the infected cells and they were cultured overnight as described above. The next day, the supernatant was harvested and clarified by centrifugation at 300g for 10 min and aliquots stored at −80 °C.

Neutralization assays were performed by incubating pseudoviruses with serial dilutions antibodies, and scored by the reduction in luciferase gene expression. In brief, Vero E6 cells were seeded in a 96-well plate at a concentration of 2 × 10^4^ cells per well. Pseudoviruses were incubated the next day with serial dilutions of the test samples in triplicate for 30 mins at 37 °C. The mixture was added to cultured cells and incubated for an additional 24 hrs. The luminescence was measured by Britelite plus Reporter Gene Assay System (PerkinElmer). IC_50_ was defined as the dilution at which the relative light units were reduced by 50% compared with the virus control wells (virus + cells) after subtraction of the background in the control groups with cells only. The IC_50_ values were calculated using non-linear regression in GraphPad Prism.

### Antibody gene assignments and genetic analyses

The 17 SARS-COV-2 neutralizing antibodies were collected from seven publications. We annotated these antibodies using IgBLAST-1.16.0 with the default parameters (Ye et al., 2013). For antibodies which have cDNA sequences deposited, the V and J genes were assigned using SONAR version 2.0 (https://github.com/scharch/sonar/) with germline gene database from IMGT (Lefranc, 2008; Schramm et al., 2016). For each antibody, the N-addition, D gene, and P-addition regions were annotated by IMGT V-QUEST (Brochet et al., 2008). To identify somatic hypermutations, each antibody sequence was aligned to the assigned germline gene using MUSCLE v3.8.31 (Edgar, 2004). Somatic hypermutations were identified from the alignment. In addition, the analysis of single cell antibody repertoire sequencing data of SARS-CoV-2 patient 2 from (Liu et al., 2020a), showed that 29 of the 38 unique transcripts assigned to IGLV2-14*01 share nucleotide mutations G156T and T165G. These mutations lead to amino acid mutations E50D and N53K. Both nucleotide mutations are also observed in 82 of 90 unique IGLV2-14 transcripts from patient 1 of the same study. Because these transcripts having different VJ recombination and paired with different heavy chain genes, the chances that the two convergent mutations are the results of somatic hypermutation are very low. Thus, we suspect that both donors contain a new IGLV2-14 gene allele (IGLV2-14*0X), which was deposited to European Nucleotide Archive (ENA) with project accession numbers: PRJEB31020. Light chain of 1-87 was assigned to the IGLV2-14*0X allele.

### Cryo-EM Samples Preparation

Samples for cryo-EM grids preparation were produced by mixing purified SARS-CoV-2 S2P spike (final trimer concentration of 0.33 mg/mL) with NTD-directed Fabs in a 1:9 molar ratio, followed by incubation on ice for 1 hr. The final buffer for 1-87, 4-18 and 5-24 complexes was 10 mM sodium acetate, 150 mM NaCl, pH 4.5; the final buffer for 1-68, 2-17, 2-51 and 4-8 complexes was 10 mM sodium acetate, 150 mM NaCl, pH 5.5. n-Dodecyl β-D-maltoside (DDM) at a final concentration of 0.005% (w/v) was added to the mixtures to prevent aggregation during vitrification. Cryo-EM grids were prepared by applying 2 µL of sample to a freshly glow-discharged carbon-coated copper grid (CF 1.2/1.3 300 mesh); the sample was vitrified in liquid ethane using a Vitrobot Mark IV with a wait time of 30 s and a blot time of 3 s.

### Cryo-EM Data Collection, Processing and Structure Refinement

Cryo-EM data were collected using the Leginon software (Suloway et al., 2005) installed on a Titan Krios electron microscope operating at 300 kV, equipped with a Gatan K3-BioQuantum direct detection device. The total dose was fractionated for 3 s over 60 raw frames or 2 s over 40 raw frames. Motion correction, CTF estimation, particle extraction, 2D classification, ab initio model generation, 3D refinements and local resolution estimation for all datasets were carried out in cryoSPARC 2.15 (Punjani et al., 2017); particles were picked using Topaz (Bepler et al., 2019). Symmetry expansion and focused classification in RELION 3.1 (Scheres, 2012) was used for S2P spike complex with 2-51. The interface between NTD and the Fab was locally refined by using a mask that included NTD and the variable domains of the Fab; symmetry-expanded particles in C3 were used in the local refinement for S2P spike complexes with 4-18 and 5-24. The 4-8 interface was locally refined following particle subtraction without symmetry expansion using a mask over the same region. The density at the interface was well-defined for S2P spike complexes with 1-87, 4-8, 4-18 and 5-24, providing structural details of antibody binding to NTD.

SARS CoV-2 S2P spike density was modeled using PDB entry 6VXX (Walls et al., 2020), as initial template. The RBDs and were initially modeled using PDB entry 7BZ5 (Wu et al., 2020); the NTDs were initially modeled using PDB entry 6ZGE (Wrobel et al., 2020). The initial models for all Fab variable regions were obtained using the SAbPred server (Dunbar et al., 2016).

Automated and manual model building were iteratively performed using real space refinement in Phenix (Adams et al., 2004) and Coot (Emsley and Cowtan, 2004) respectively. Geometry validation and structure quality assessment were performed using EMRinger (Barad et al., 2015) and Molprobity (Davis et al., 2004). Map-fitting cross correlation (Fit-in-Map tool) and figures preparation were carried out using PyMOL and UCSF Chimera (Pettersen et al., 2004) and Chimera X (Pettersen et al., 2021). A summary of the cryo-EM data collection, reconstruction and refinement statistics is shown in Table S1.

### X-ray Crystallography Sample Preparation, Data Collection, Structure Solution and Refinement

Purified SARS-CoV-2 spike N-terminal domain (NTD) and 2-51 Fab were mixed at a 1:1 molar ratio and incubated at 4 °C for 1 hr; the Fab-NTD complex was purified by SEC on a Superdex 200 column in buffer 10 mM Tris, 150 mM NaCl, pH 7.4. Fractions containing the complex were combined and concentrated to a total protein concentration of 7.5 mg/mL for crystal screening by the sitting drop vapor diffusion method at 25 °C. Diffracting crystals of NTD in complex with 2-51 Fab grew in 0.16 M calcium acetate, 0.08 M sodium cacodylate, 14.4% PEG 8000, 20% glycerol, pH 6.5. For data collection, crystals were cryo-protected by briefly soaking in reservoir solution supplemented with 35% (v/v) glycerol before flash-cooling in liquid nitrogen. X-ray diffraction data was collected to 3.44 Å resolution at 100 K from a single flash-cooled crystal on beamline 24ID-C at the Advance Photon Source (APS) at Argonne National Laboratory. Diffraction data were processed with XDS (Kabsch, 2010) and scaled using AIMLESS (Evans and Murshudov, 2013) from the CCP4 software suite (Winn et al., 2011). Molecular replacement was performed with Phaser (McCoy, 2007) using the structure of NTD in complex with 4A8 (extracted from the whole spike-Fab structure from PDB entry 7C2L) as search model. Structure refinement was performed with a 3.65 Å high-resolution cutoff using Phenix refine (Adams et al., 2010) and PDB-redo (Joosten et al., 2014) alternated with manual model building using Coot. The Molprobity server was used for geometry validation and structure quality assessment. A summary of the X-ray crystallography data collection and refinement statistics is shown in Table S2.

### Calculation of antibody angle of approach

The angles of antibody approach to the NTD of SARS-CoV-2 spike were calculated with UCSF Chimera (Pettersen et al., 2014). To define the latitudinal and longitudinal access of an antibody to the viral spike, we first defined two reference axes: the 3-fold axis of the spike trimer and a line perpendicular to the 3-fold axis and passing through the Cα atom of Trp104 located in the hydrophobic core of the NTD. We then defined the axis of antibody as the long axis of the Fab. The latitudinal access, which describes the freedom between the viral and host cell membranes, was defined as the angle between the antibody axis and the 3-fold axis; the longitudinal access, which describes the freedom within the plane of the membrane, was defined as the angle between the antibody axis and the other reference axis. The angles between two axes were calculated with the built-in function of UCSF Chimera, and expressed in 0-to-180-degree scale for the latitudinal angles and 0-to-360-degree scale counter-clockwise to show the longitudinal angles. To compare the relative orientations of antibody heavy and light chains, we used a vector going from the center of heavy chain variable domain to that of the light chain variable domain.

### Glycan analysis

To estimate the effect of glycan shield on protein surface residue, we used an in-house algorithm to quantify the number of glycan atoms associated with each residue. Briefly, the number of glycan atoms per protein surface residue was counted within 34 Å radius distance cutoff, while the glycans do not provide the shielding effect were excluded in the calculation. This analysis was performed on each trajectory of molecular dynamics simulation (Casalino et al., 2020) to average the glycan atom counts for each residue.

### Interface definition and net charge computation

The residues of each antibody paratope were obtained by running PDBePISA (Krissinel and Henrick, 2007), with the default parameters. The hydrophobic interaction residues among antibodies were defined by the buried surface area (BSA) that large than 20 Å^2^. The antigen contact region for germline genes were adopt from (Sela-Culang et al., 2013), which includes additional interactions not accounted for in the CDRs. The number of charged residues were counted and the summation of their charges was used to quantify the net charge of selected residues.

### C α distance calculation

The Cα distances per residue between superimposed ligand-free and antibody-bound NTD were calculated by Python 3.8. The 8 NTD-antibody complexes (NTD-5-24, NTD-4-8, NTD-2-17, NTD-4-18, NTD-1-87, NTD-4A8, NTD-2-51, NTD-1-68) were used as antibody bound structures, and the averaged Cα distances per residue of 8 complexes were analyzed.

### RMSD calculation

The root mean square deviation (RMSD) of antibody epitopes was calculated by PyMOL 2.4 between two superimposed NTDs – ligand-free and antibody-bound structures. The identical ligand-free and antibody-bound NTDs described in Cα distance analysis were used in the calculation. Only matching atoms with the same name and number in both epitopes were included in the calculation. The epitope residues of antibodies were obtained by running FreeSASA (Mitternacht, 2016) with probe radius 1.4 Å. The residues with non-zero surface area difference between NTD and NTD-antibody complex structures were selected as protein epitope residues to the corresponding antibodies.

### Sequence entropy of betacoronavirus spike

Human coronavirus reference amino acid sequences of OC43 (UniProt ID: P36334), HKU1 (UniProt ID: Q5MQD0), SARS (UniProt ID: P59594), and MERS (UniProt ID: W5ZZF5), as well as the initial SARS-CoV-2 (GenBank: QHO60594.1) sequence reported in Washington state were aligned using MAFFT software with default parameters (Katoh et al., 2002). Subsequently, we used the R bio3d package’s function Conserv with default parameters to estimate sequence conservation at all alignment position

### QUANTIFICATION AND STATISTICAL ANALYSIS

The statistical analyses for the pseudovirus neutralization assessments were performed using GraphPad Prism. The SPR data were fitted using Biacore Evaluation Software. Cryo-EM data were processed and analyzed using cryoSPARC and Relion. Cryo-EM and crystallographic structural statistics were analyzed using Phenix, Molprobity, EMringer and Chimera. The correlations were performed in R. Statistical details of experiments are described in Method Details or figure legends.

