## Supplemental Figures and Tables for "Potent SARS-CoV-2 Neutralizing Antibodies Directed Against Spike N-Terminal Domain Target a Single Supersite"

**A**

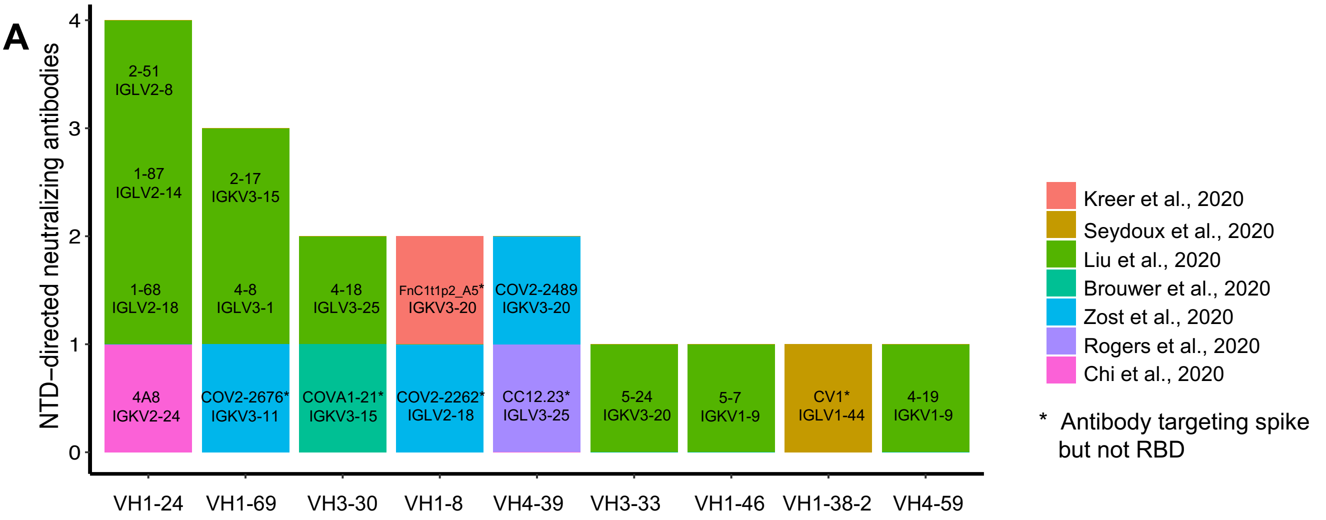


**B**

| mAb | HV | LV | Neutralization IC50 (µg/mL) | Epitope | Reference |
|---|---|---|---|---|---|
| COV2-2489 | IGHV4-39 | IGKV3-20 | 16. | NTD | Zost et al., 2020 |
| COV2-2262 | IGHV1-8 | IGLV2-18 | 6.0 | S, Non-RBD | Zost et al., 2020 |
| COV2-2676 | IGHV1-69 | IGKV3-11 | 9.6 | S, Non-RBD | Zost et al., 2020 |
| FnC1t1p2_A5 | IGHV1-8 | IGKV3-20 | 16. | S, Non-RBD | Kreer et al., 2020 |
| 4A8 | IGHV1-24 | IGKV2-24 | 0.39 | NTD | Chi et al., 2020 |
| COVA1-21 | IGHV3-30 | IGKV3-15 | 0.16 | S, Non-RBD | Brouwer et al., 2020 |
| CC12.23 | IGHV4-39 | IGLV3-25 | 8.7 | S, Non-RBD | |
| CV1 | IGHV4-38-2 | IGLV1-44 | 15. | S, Non-RBD | Seydoux et al., 2020 |
| 4-8 | IGHV1-69 | IGLV3-1 | 0.009 | NTD | Liu et al., 2020 |
| 1-68 | IGHV1-24 | IGLV2-18 | 0.014 | NTD | Liu et al., 2020 |
| 1-87 | IGHV1-24 | IGLV2-14 | 0.086 | NTD | Liu et al., 2020 |
| 2-51 | IGHV1-24 | IGLV2-8 | 0.007 | NTD | Liu et al., 2020 |
| 5-24 | IGHV3-33 | IGKV3-20 | 0.008 | NTD | Liu et al., 2020 |
| 4-19 | IGHV4-59 | IGKV1-9 | 0.109 | NTD | Liu et al., 2020 |
| 4-18 | IGHV3-30 | IGLV3-25 | 0.020 | NTD | Liu et al., 2020 |
| 5-7 | IGHV1-46 | IGKV1-9 | 0.033 | NTD | Liu et al., 2020 |
| 2-17 | IGHV1-69 | IGKV3-15 | 0.007 | NTD | Liu et al., 2020 |

**C**

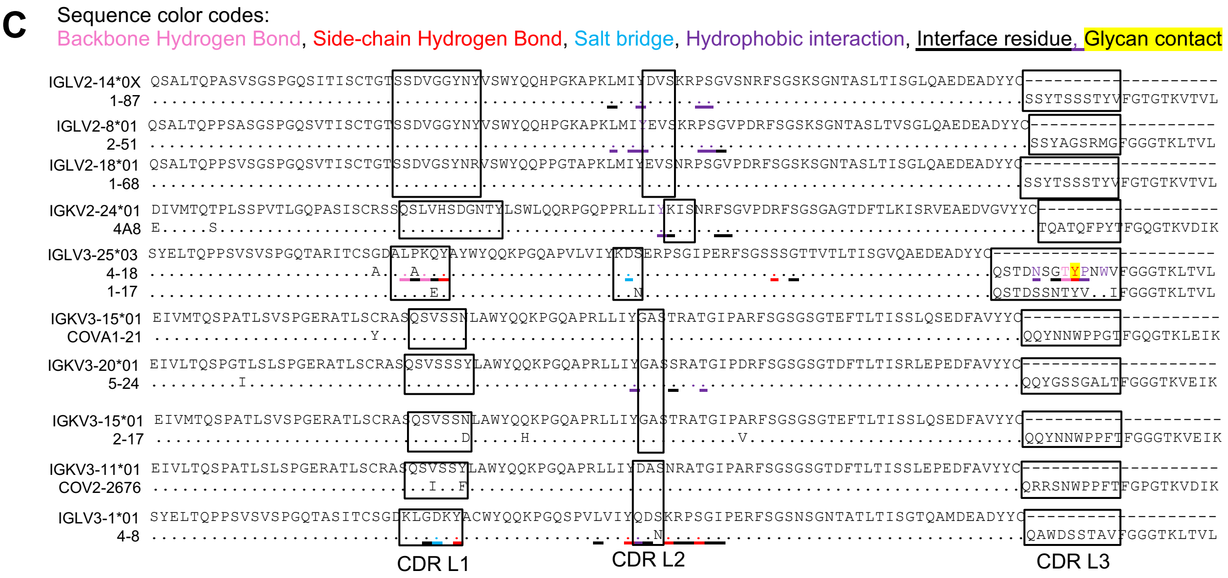


**Figure S1. NTD-directed neutralizing antibodies isolated from convalescent donors show enrichment of four VH genes, Related to Figures 1-3.**

(A) Published SARS-CoV-2 neutralizing antibodies targeting NTD or non-RBD spike epitopes.

(B) Table of published NTD antibodies with neutralization data.

(C) Light chain sequence alignment for VH1-24-derived antibodies, for VH3-30/33-derived antibodies, and for VH1-69-derived antibodies. 1-87 was assigned to a novel germline gene IGLV1-24*0X.

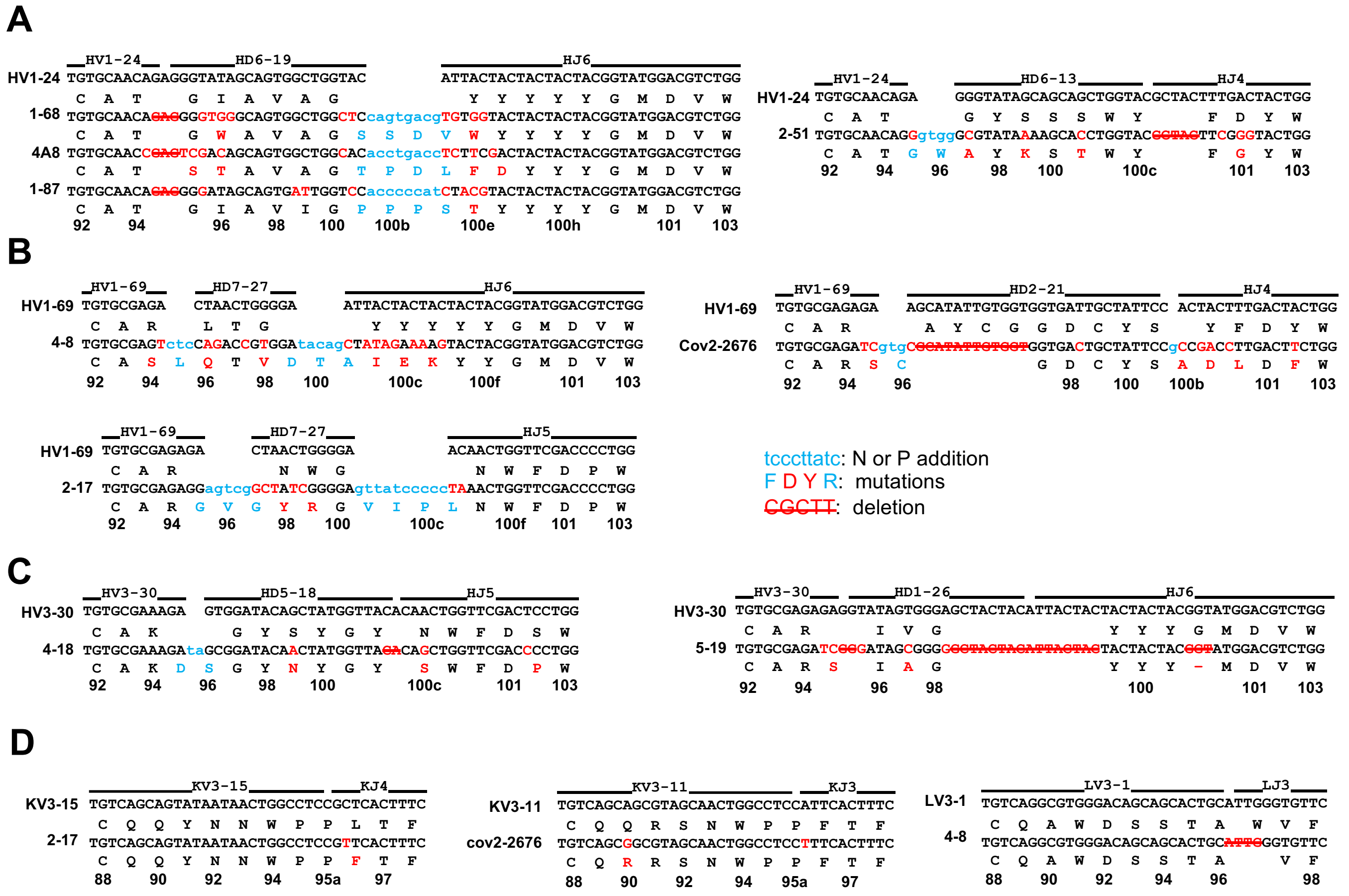


**Figure S2. Heavy chain VDJ junction analysis for NTD neutralizing antibodies and light chain VJ junction analysis for antibodies derived from VH1-69, Related to Figures 1-3.**
Germline nucleotide and amino acid residues are shown in black with the corresponding junctions colored in light blue. Somatic hypermutations are colored in red. Nucleotides deleted by exonuclease trimming are indicated with strikethrough. The lower-case blue nucleotides represent the N and P nucleotide additions at the junctions.

(A) VH1-24-derived antibodies.
(B) VH1-69-derived antibodies.
(C) VH3-30/33-derived antibodies.
(D) Light chain VH junctional analysis for VH1-69-derived antibodies.

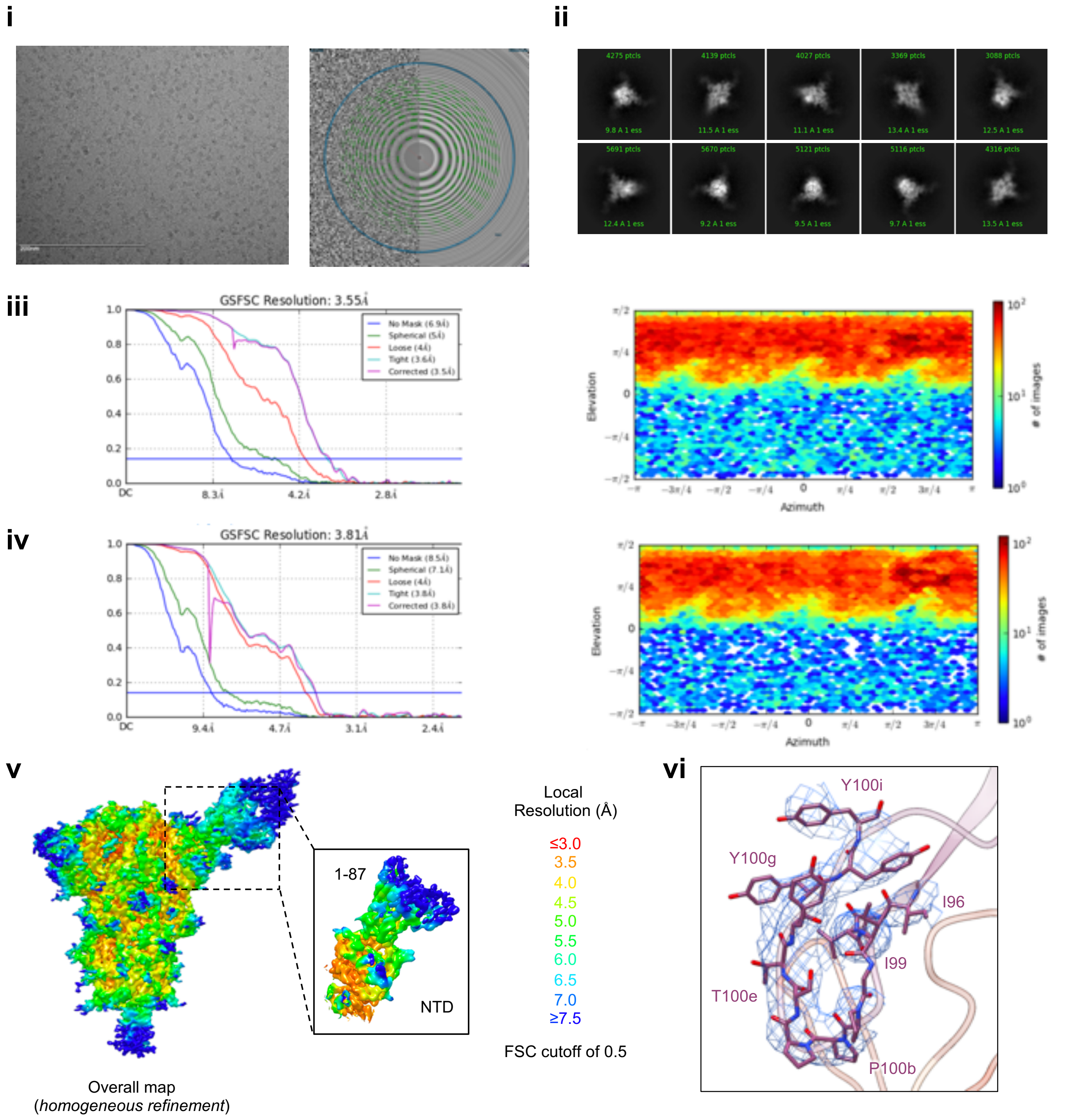


**Figure S3A. Cryo-EM details of 1-87 Fab in complex with SARS-CoV-2 S2P spike, Related to Figure 1.**

(i) Representative micrograph and CTF of the micrograph are shown.

(ii) Representative 2D class averages are shown.

(iii) The gold-standard Fourier shell correlation resulted in a resolution of 3.55 Å for the overall map using non-uniform refinement (left panel); the orientations of all particles used in the final refinement are shown as a heatmap (right panel); the observed preferred orientation did not preclude the generation of a high-quality map.

(iv) The gold-standard Fourier shell correlation resulted in a resolution of 3.81 Å for the masked local refinement of the NTD:1-87 interface (left panel); the orientations of all particles used in the local refinement are shown as a heatmap (right panel). the observed preferred orientation did not preclude the generation of a high-quality map.

(v) The local resolution of the final overall map and locally refined map are shown, generated through cryoSPARC using an FSC cutoff of 0.5.

(vi) Representative density is shown for the CDR H3 loop of 1-87 contacting NTD. CDR H3 carbon atoms are colored in magenta, oxygen in red, nitrogen in blue; NTD is colored in orange.

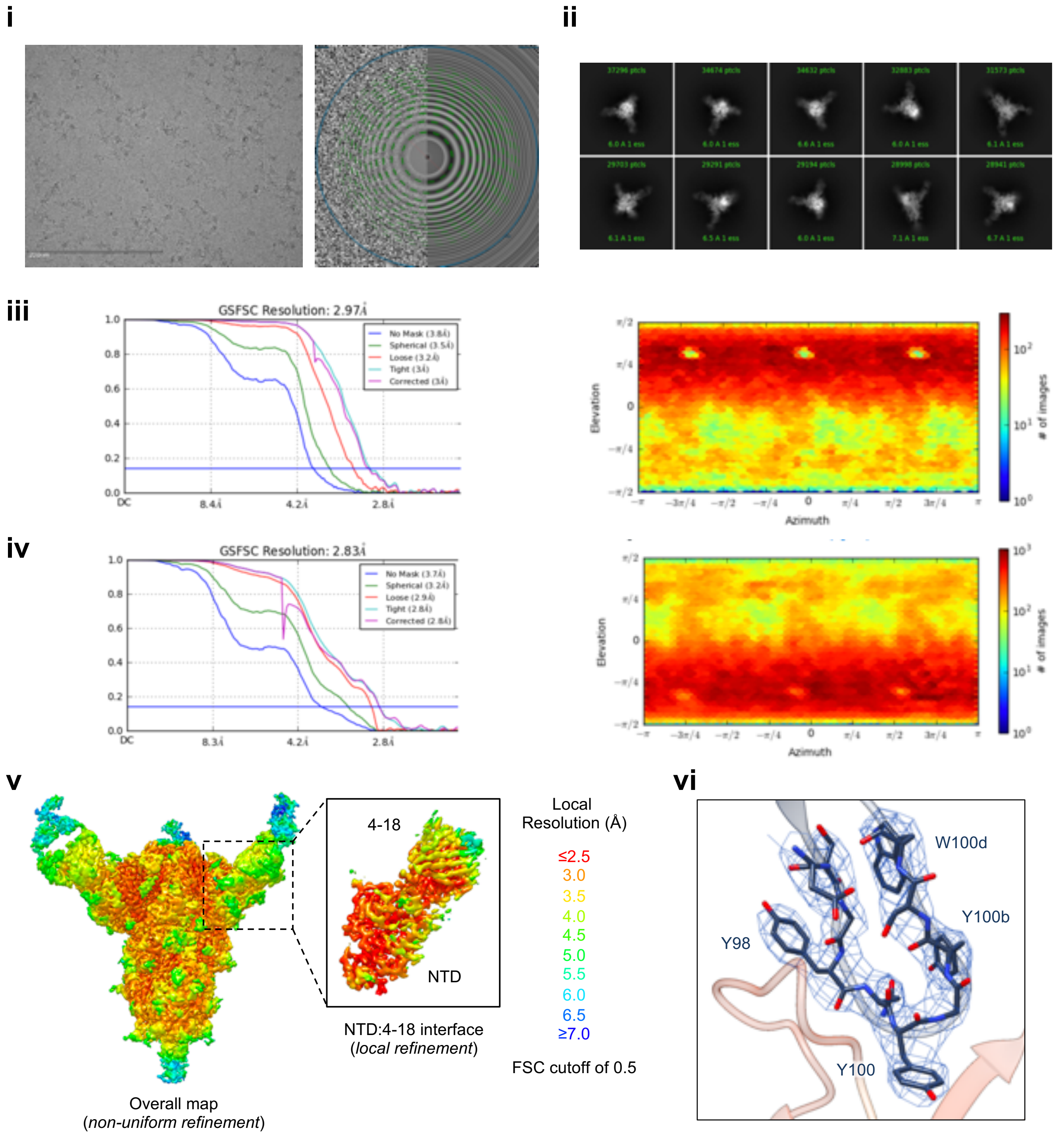


**Figure S3B. Cryo-EM details of 4-18 Fab in complex with SARS-CoV-2 S2P spike, Related to Figure 2.**
(i) Representative micrograph and CTF of the micrograph are shown.
(ii) Representative 2D class averages are shown.
(iii) The gold-standard Fourier shell correlation resulted in a resolution of 2.97 Å for the overall map using non-uniform refinement (left panel); the orientations of all particles used in the final refinement are shown as a heatmap (right panel).
(iv) The gold-standard Fourier shell correlation resulted in a resolution of 2.83 Å for the masked local refinement of the NTD:4-18 interface (left panel) obtained using symmetry expansion in C3; the orientations of all particles used in the local refinement are shown as a heatmap (right panel).
(v) The local resolution of the final overall map and locally refined map are shown, generated through cryoSPARC using an FSC cutoff of 0.5.
(vi) Representative density is shown for the CDR H3 loop of 4-18 contacting NTD. CDR H3 carbon atoms are colored in dark blue, oxygen in red, nitrogen in blue; NTD is colored in orange.

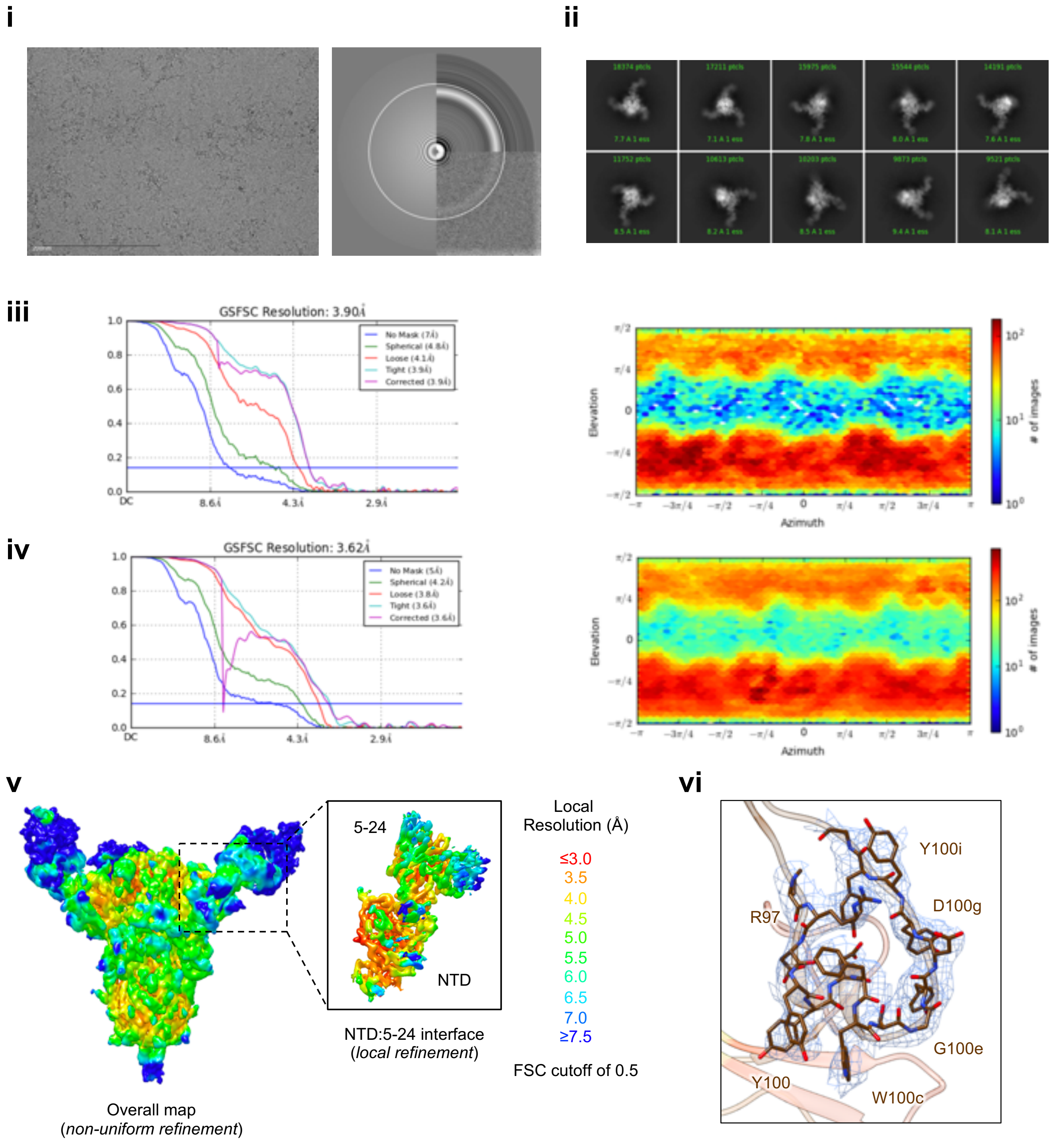


**Figure S3C. Cryo-EM details of 5-24 Fab in complex with SARS-CoV-2 S2P spike, Related to Figure 2.**
(i) Representative micrograph and CTF of the micrograph are shown.
(ii) Representative 2D class averages are shown.
(iii) The gold-standard Fourier shell correlation resulted in a resolution of 3.90 Å for the overall map using non-uniform refinement (left panel); the orientations of all particles used in the final refinement are shown as a heatmap (right panel).
(iv) The gold-standard Fourier shell correlation resulted in a resolution of 3.62 Å for the masked local refinement of the NTD:5-24 interface (left panel) obtained using symmetry expansion in C3; the orientations of all particles used in the local refinement are shown as a heatmap (right panel).
(v) The local resolution of the final overall map and locally refined map are shown, generated through cryoSPARC using an FSC cutoff of 0.5.
(vi) Representative density is shown for the CDR H3 loop of 5-24 contacting NTD. CDR H3 carbon atoms are colored in brown, oxygen in red, nitrogen in blue; NTD is colored in orange.

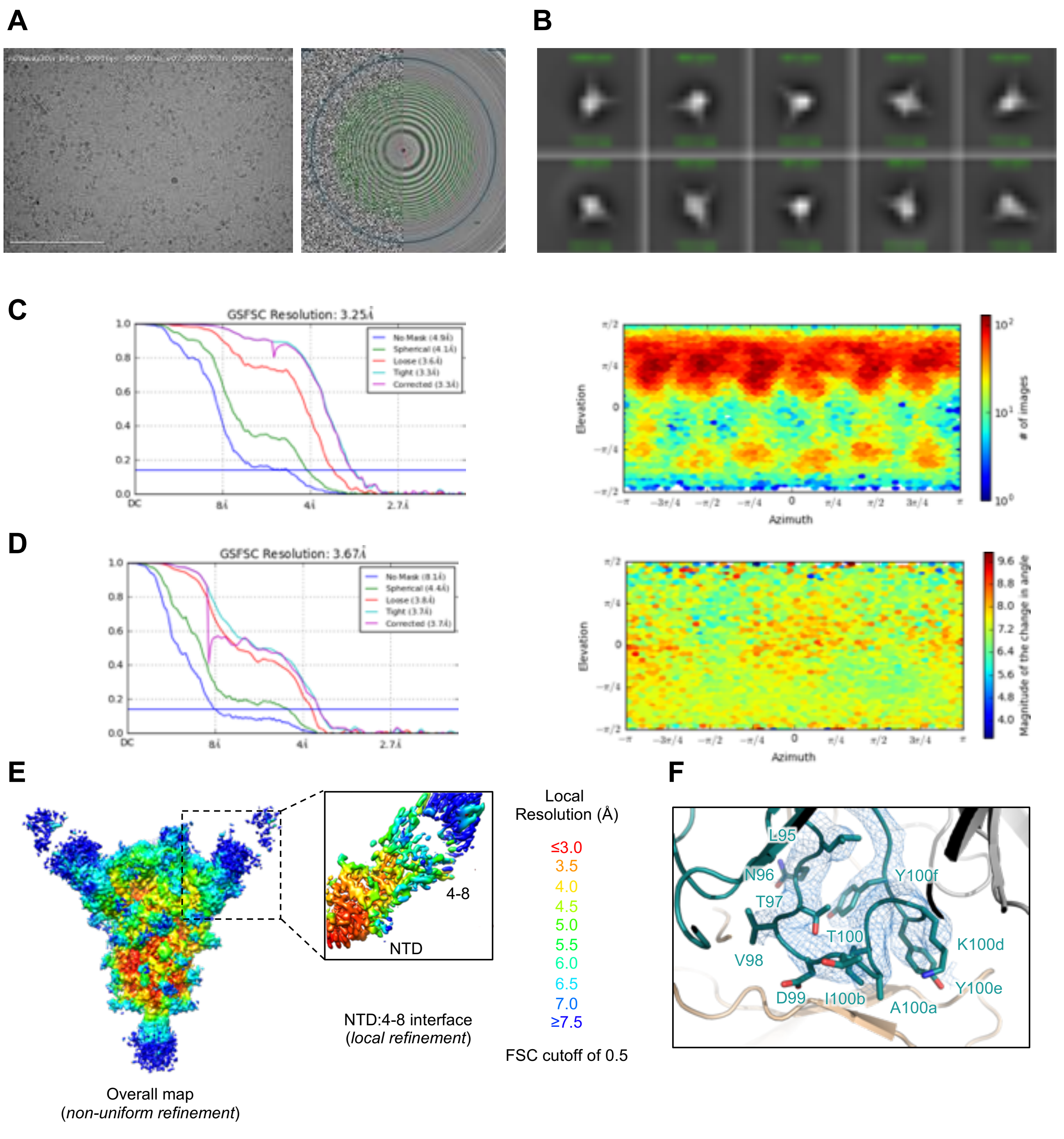


**Figure S3D. Cryo-EM details of 4-8 Fab in complex with SARS-CoV-2 S2P spike, Related to Figure 3.**
(i) Representative micrograph and CTF of the micrograph are shown.
(ii) Representative 2D class averages are shown.
(iii) The gold-standard Fourier shell correlation resulted in a resolution of 3.25 Å for the overall map using non-uniform refinement with C1 symmetry (left panel); the orientations of all particles used in the final refinement are shown as a heatmap (right panel).
(iv) The gold-standard Fourier shell correlation resulted in a resolution of 3.67 Å for the masked local refinement of the NTD:4-8 interface (left panel) obtained using particle subtraction followed by local refinement; the orientations of all particles used in the local refinement are shown as a heatmap (right panel).
(v) The local resolution of the final overall map and locally refined map are shown, generated through cryoSPARC using an FSC cutoff of 0.5.
(vi) Representative density is shown for the CDR H3 loop of 4-8 contacting NTD. CDR H3 carbon atoms are colored in dark teal, oxygen in red, nitrogen in blue; NTD is colored in orange.

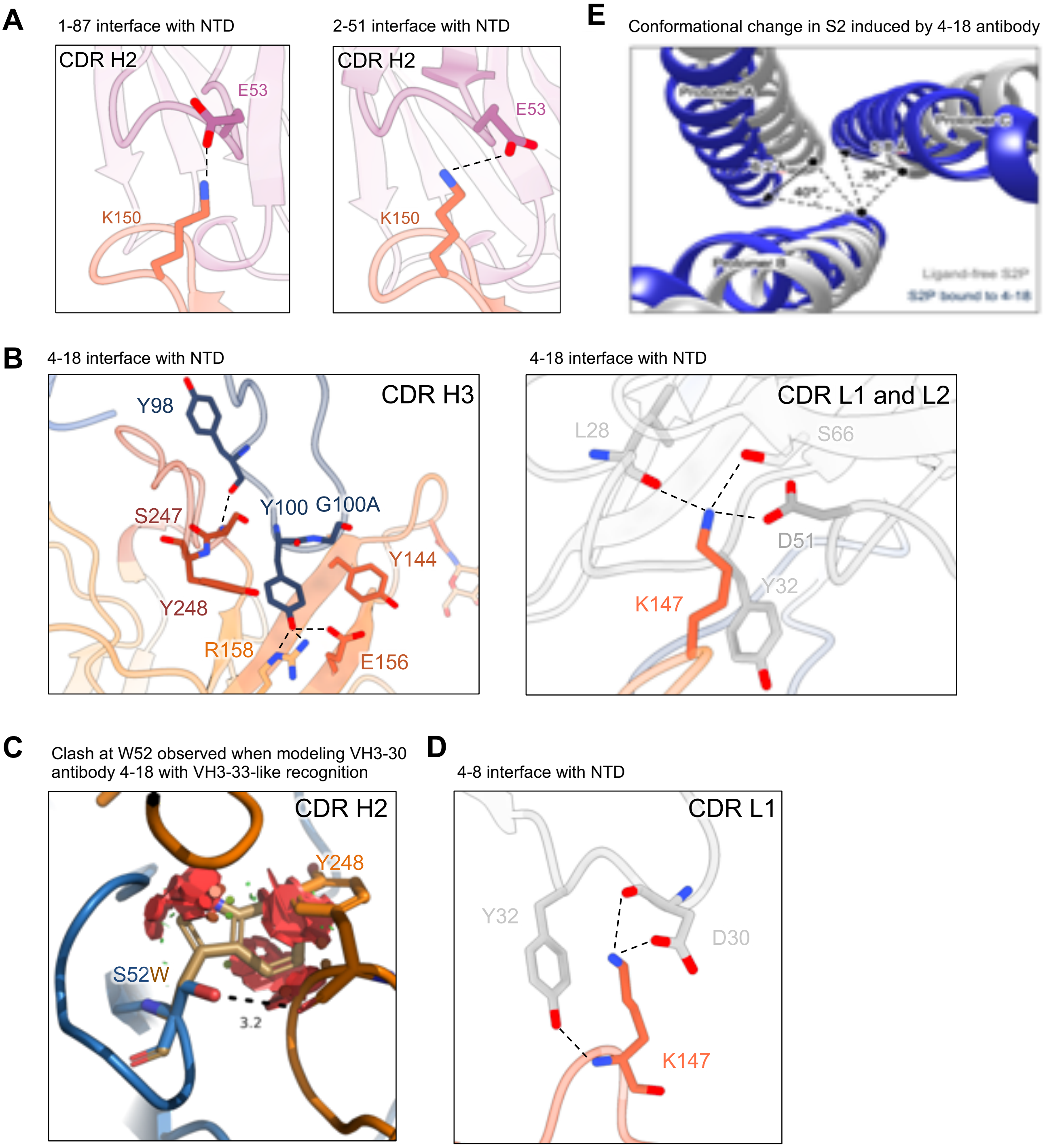


**Figure S4. Additional observations from 1-87, 4-18 and 4-8 complexes, Related to Figures 1-3.**

(A) The main interaction observed in CDR H2 for VH1-24-derived antibodies is a salt bridge between Glu53 and Lys150, observed in both 1-87 (left panel) and 2-51 (right panel). NTD is colored in orange; CDHR H2 is colored in magenta. Nitrogen atoms are colored in blue, oxygen atoms in red; hydrogen bonds are represented as dashed lines.

(B) Expanded view of 4-18 interactions with NTD showing recognition in CDR H3 (left panel), and recognition in CDR L1 and L2 (right panel). NTD regions N3 (residues 141-156) and N5 (residues 246-260) are colored in shades of orange; CDR H3 is colored in dark blue; CDR L1 and L2 are colored in shades of gray.

(C) The S52W substitution between VH3-30 and VH3-33 is incompatible with the binding mode of 4-18. Mutating Ser52 (blue) to Tryptophane (brown) would bring major steric clashes (red plate) between 4-18 heavy chain (blue) and NTD (orange). The hydrogen bond between Ser52 and Tyr248 on NTD is represented as a dashed line.

(D) Expanded view of 4-8 interactions with NTD showing recognition in CDR L1, colored as in (B)

(E) Conformational change in the S2 region of spike induced by 4-18 antibody binding.

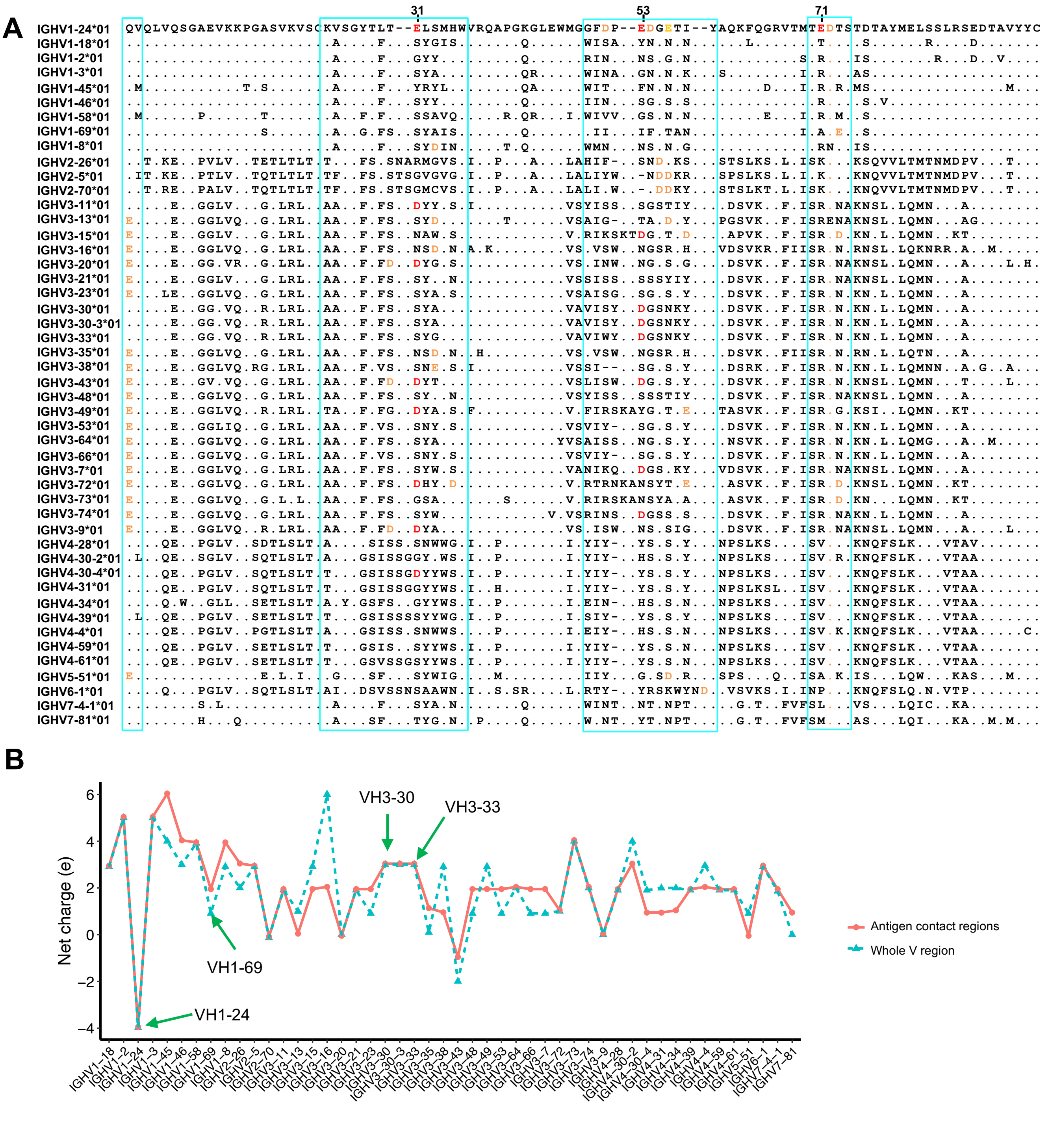


**Figure S5. VH1-24 is the most negatively charged germline gene, Related to Figure 1.**

(A) Multiple sequence alignment of the *01 allele for all VH genes. The cyan boxes show antigen contact regions defined by Sela-Culang et al. (2013), which include additional interactions not accounted for in the CDRs. The dots represent conserved residues compared with the VH1-24 gene. The negative charge residues at Kabat position 31, 53 and 71 are colored in red, and other negative charge residues within antigen contact regions are colored in orange.

(B) Net charge distribution of all VH genes. The cyan triangles represent the net charge of the whole V region, the red dots represent the net charge in antigen contact regions in panel A. The green arrows highlight the VH1-24, VH1-69, VH3-30 and VH3-33 germline genes.

**A**

| Antibody | ka ($M^{-1}s^{-1}$) | kd ($s^{-1}$) | $K_D^{app}$ (nM) |
|---|---|---|---|
| 1-87 | 114444 | 6.1775E-06 | 0.1359 |
| 1-87 IH30T | 80919 | 6.1775E-06 | 0.0763 |
| 1-87 AH55G | 65323 | 1.2732E-04 | 1.9490 |
| 1-87 IH30T AH55G | 52927 | 1.1720E-04 | 2.2145 |
| 1-68 | 99785 | 2.4129E-04 | 2.4181 |
| 1-68 IH30T | 71112 | 7.4050E-04 | 10.4131 |
| 1-68 AH55G | 75649 | 4.2248E-04 | 5.5848 |
| 1-68 IH30T AH55G | 78662 | 3.1756E-03 | 40.3702 |
| 2-51 | 149981 | 5.4046E-04 | 3.6035 |
| 2-51 IH30T | 20299 | 2.9341E-03 | 14.4540 |
| 2-51 VH55G | 230715 | 3.7759E-03 | 16.3660 |
| 2-51 IH30T VH55G | 367225 | 2.6717E-02 | 72.7536 |
| 4-18 | 34869 | 1.4859E-07 | 0.0043 |
| 4-18 HH58Y AL29P TL91A NL93S | 42767 | 1.7901E-03 | 41.8578 |
| 5-24 | 106110 | 4.4160E-05 | 0.4162 |
| 5-24 LH27F KH56N | 97333 | 4.2056E-04 | 4.3208 |
| 2-17 | 36080 | 1.1175E-08 | 0.0003 |
| 2-17 NH51I DL32N | 106299 | 7.5238E-04 | 7.0779 |
| 4-8 | 125519 | 5.6236E-04 | 4.4802 |
| 4-8 HH32Y TH35S NL52S | 127990 | 2.6868E-03 | 20.9922 |
| 4A8 | 355683 | 5.0827E-04 | 1.4290 |

**B**

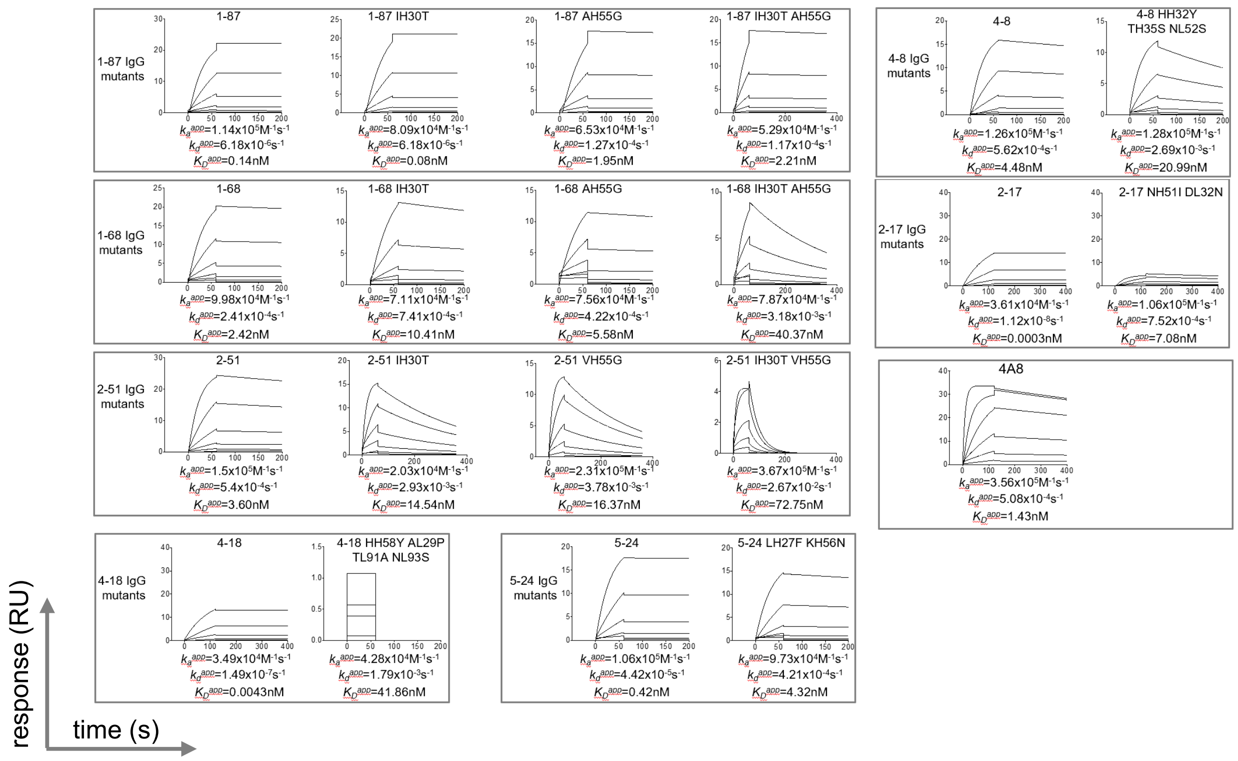


**C**

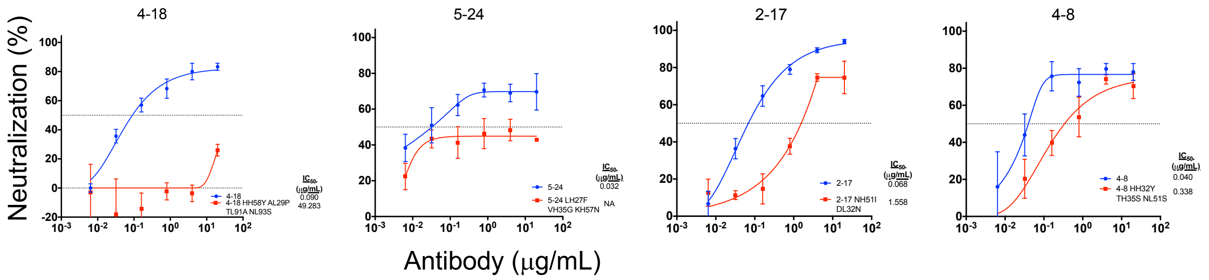


**Figure S6. Effects of somatic hypermutation on binding affinity and neutralization potency of NTD antibodies, Related to Figures 1, 2, 3 and 6.**

(A) Apparent SARS-CoV-2 spike binding affinity of NTD-directed antibodies (IgGs) show that somatic hypermutations significantly improve binding affinity.

(B) Surface plasmon resonance profiles of NTD-directed antibodies and revertants.

(C) Pseudovirus neutralization profiles of NTD-directed antibodies show that somatic hypermutations significantly improve neutralization potency. Wildtype antibodies are blue and mutants are red. Mean±SEM is shown for each data point.

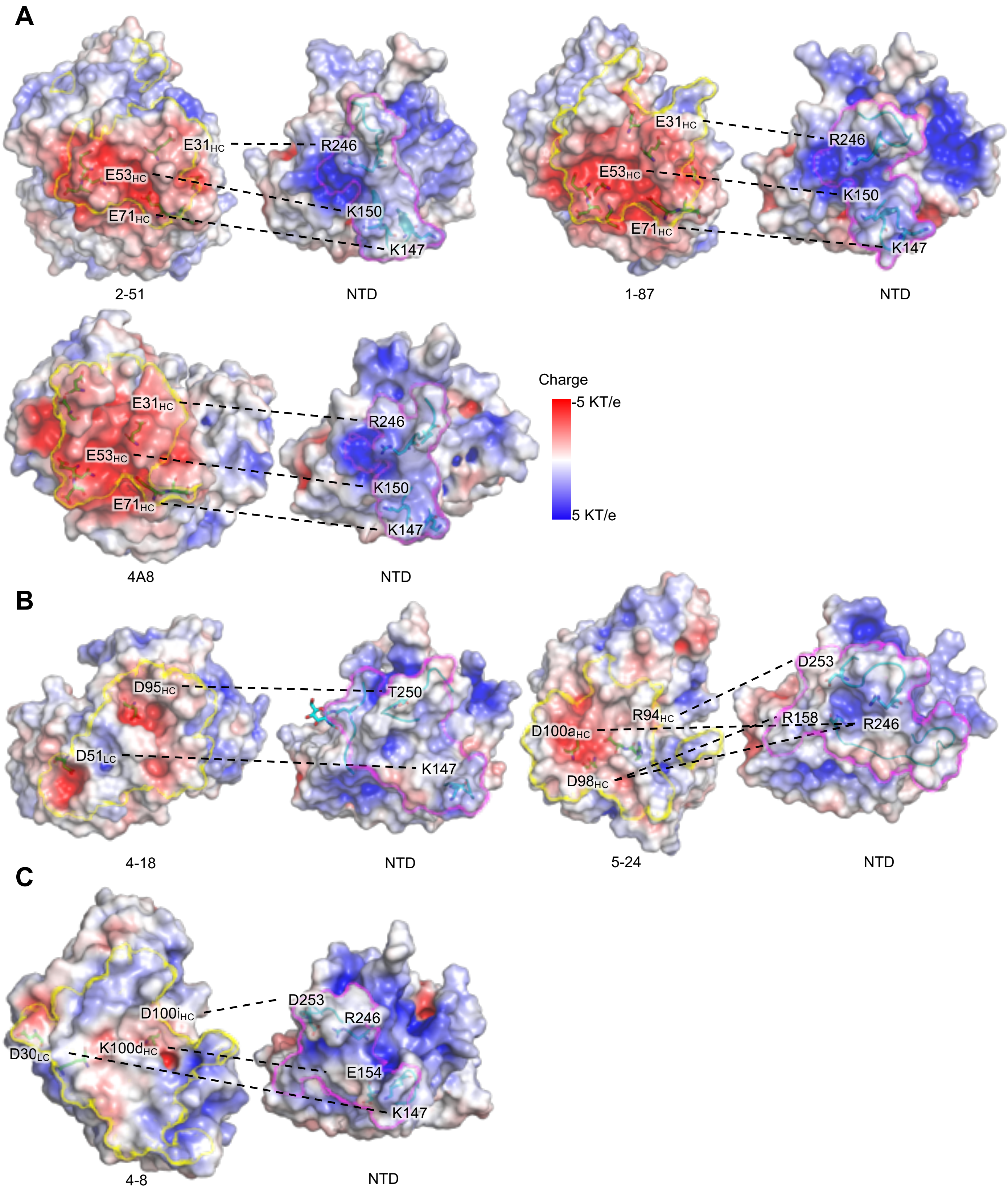


**Figure S7. NTD-directed neutralizing antibodies are electronegative and target the electropositive supersite, Related to Figures 1, 2, 3 and 6.**

(A) Electrostatic potential for VH1-24-derived antibodies. The blue surface shows electropositive charge potential, and the red shows negative charge potential. The paratope and epitope are highlighted by yellow and magenta boundaries, respectively. Residues involved in charge-charge interactions between antibody and NTD are linked by dashed lines.

(B) Electrostatic potential for VH3-30 (4-18) and VH3-33 (5-24) derived antibodies.

(C) Electrostatic potential for VH1-69-derived antibodies.

**Table S1. Cryo-EM Data Collection and Refinement Statistics, Related to Figures 1-3.**

| SARS-CoV-2 S2P complex | 1-87 Fab | 4-18 Fab | 5-24 Fab | 4-8 Fab | 1-68 Fab | 2-51 Fab | 2-17 Fab |
|---|---|---|---|---|---|---|---|
| **EMDB ID** | EMD-23125 | EMD-23126 | EMD-23127 | EMD-XXXXX | EMD-23150 | EMD-23151 | EMD-XXXXX |
| **PDB ID** | 7L2D | 7L2E | 7L2F | XXXX | | | XXXX |
| Data Collection | | | | | | | |
| Microscope | FEI Titan Krios | FEI Titan Krios | FEI Titan Krios | FEI Titan Krios | FEI Titan Krios | FEI Titan Krios | FEI Titan Krios |
| Voltage (kV) | 300 | 300 | 300 | 300 | 300 | 300 | 300 |
| Electron dose ($e^-$/Å$^2$) | 41.92 | 41.92 | 41.92 | 52.56 | 41.92 | 41.92 | 51.69 |
| Detector | Gatan K3 BioQuantum | Gatan K3 BioQuantum | Gatan K3 BioQuantum | Gatan K3 BioQuantum | Gatan K3 BioQuantum | Gatan K3 BioQuantum | Gatan K3 BioQuantum |
| Pixel Size (Å) | 1.07 | 1.07 | 1.07 | 1.058 | 1.07 | 1.07 | 1.058 |
| Defocus Range (µm) | -0.8/-2.5 | -0.8/-2.5 | -0.8/-2.5 | -0.1/-3.6 | -0.8/-2.5 | -0.8/-2.5 | -0.3/-3.9 |
| Magnification | 81000 | 81000 | 81000 | 81000 | 81000 | 81000 | 81000 |
| Reconstruction | | | | | | | |
| Software | cryoSPARC v2.15 | cryoSPARC v2.15 | cryoSPARC v2.15 | cryoSPARC v2.15 | cryoSPARC v2.15 | cryoSPARC v2.15 | cryoSPARC v2.15 |
| Particles | 62,479 | 280,327 | 115,545 | 88,375 | 34,450 | 190,557 | 43,224 |
| Symmetry | C1 | C3 | C1 | C1 | C1 | C1 | C3 |
| Box size (pix) | 390 | 392 | 400 | 380 | 440 | 384 | 384 |
| Resolution (Å) ($FSC_{0.143}$) | 3.55 | 2.97 | 3.90 | 3.25 | 3.80 | 3.71 | 4.03 |
| Refinement | | | | | | | |
| Software | Phenix 1.18 | Phenix 1.18 | Phenix 1.18 | Phenix 1.18 | | | |
| Protein residues | 3492 | 4050 | 3970 | 3981 | | | |
| Chimera CC | 0.85 | 0.80 | 0.86 | 0.81 | | | |
| EMRinger Score | 2.43 | 3.18 | 1.17 | 2.07 | | | |
| R.m.s. deviations | | | | | | | |
| Bond lengths (Å) | 0.006 | 0.007 | 0.008 | 0.004 | | | |
| Bond angles (°) | 1.16 | 1.29 | 1.27 | 0.7 | | | |
| Validation | | | | | | | |
| Molprobity score | 1.41 | 1.36 | 1.41 | 1.75 | | | |
| Clash score | 4.64 | 4.30 | 4.47 | 6.1 | | | |
| Favored rotamers (%) | 100 | 100 | 100 | 100 | | | |
| Ramachandran | | | | | | | |
| Favored regions (%) | 97.0 | 97.2 | 96.9 | 94.44 | | | |
| Allowed regions (%) | 3.0 | 2.8 | 3.1 | 5.53 | | | |
| Disallowed regions (%) | 0 | 0 | 0 | 0.03 | | | |

**Table S2. X-ray Diffraction Data Collection and Refinement Statistics, Related to Figure 1.**

| | SARS-CoV-2 NTD in complex with 2-51 Fab |
|---|---|
| **PDB ID** | 7L2C |
| Data Collection | |
| Space group | $P2_1$ |
| Unit cell dimensions | |
| *a,b,c* (Å) | 66.8, 115.8, 137.6 |
| $\alpha,\beta,\gamma$ (°) | 90.0, 100.0, 90.0 |
| Resolution range (Å) | 88.03-3.44 (3.57-3.44)* |
| Total reflections | 49329 (2732) |
| Unique reflections | 25964 (1550) |
| Completeness (%) | 93.3 (47.4) |
| Redundancy | 1.9 (1.8) |
| $I/\sigma(I)$ | 2.57 (0.79) |
| $R_{merge}$ | 0.2333 (1.002) |
| $CC_{1/2}$ | 0.883 (0.322) |
| Refinement | |
| Resolution range (Å) | 88.03-3.65 |
| Number of complexes per asymmetric unit | 2 |
| $R_{work}/R_{free}$ | 21.6/27.2 |
| Number of atoms | |
| Protein | 11041 |
| Ligands | 300 |
| Water | 41 |
| B-factors | |
| Protein | 69.29 |
| Ligands | 94.04 |
| Water | 49.06 |
| R.m.s. deviations | |
| Bond lengths (Å) | 0.009 |
| Bond angles (°) | 1.48 |
| Ramachandran statistics | |
| Favored (%) | 94.11 |
| Allowed (%) | 5.89 |
| Outliers (%) | 0 |

* Values in parentheses are for the highest-resolution shell.

**Table S3. Kinetic parameters and affinities for the binding of NTD-directed antibody Fabs to SARS-CoV-2 spike, Related to Figure 6.**

| Fab | $k_a$ ($M^{-1}s^{-1}$) | $k_d$ ($s^{-1}$) | $K_D$ (nM) |
|---|---|---|---|
| **1-87** | 5.76(4)x10$^{4}$ | 1.40(6)x10$^{-4}$ | 2.44(2) |
| **1-68** | 5.69(2)x10$^{4}$ | 7.86(2)x10$^{-4}$ | 13.83(3) |
| **2-51** | 9.11(2)x10$^{4}$ | 2.67(2)x10$^{-3}$ | 29.32(3) |
| **4-18** | 1.71(1)x10$^{4}$ | 1.07(4)x10$^{-4}$ | 6.26(3) |
| **5-24** | 5.83(4)x10$^{4}$ | 6.71(6)x10$^{-5}$ | 1.15(1) |
| **2-17** | 4.0(1)x10$^{3}$ | 1.25(3)x10$^{-3}$ | 317(4) |
| **4-8** | 3.51(3)x10$^{4}$ | 2.99(2)x10$^{-3}$ | 85.0(3) |
| **4A8** | 7.88(3)x10$^{4}$ | 2.82(4)x10$^{-3}$ | 35.83(5) |
